# Prioritization of prostate cancer to immune checkpoint therapy by ranking tumors along IFN-γ axis and identification of immune resistance mechanisms

**DOI:** 10.1101/2020.10.19.345629

**Authors:** Boris Reva, Tatiana Omelchenko, Anna Calinawan, Sujit Nair, Eric Schadt, Ash Tewari

## Abstract

We propose priotirization of prostate cancer patients to PD-L1 checkpoint therapy by assessing activity of IFN-γ/PD-L1 signaling in tumor from transcriptional profile. To this end, we introduced a new approach for inferring pathway activity and suppression (IPAS) by assessing significance of positioning pathway’s genes expression levels at top (activation) or bottom (suppression) in gene expression profile of a given tumor. By ordering tumors along IFN-γ/PD-L1 axis, we determined distinct “IFN-γ-depleted” and “IFN-γ-enriched” immune subtypes, genes involved in immune evasion and potential targets for combination therapy. Using IPAS scoring method, we proposed biomarker panels for accurate ranking tumors along IFN-γ/PD-L1 axis.

## 1. Introduction

The diversity of cancer driving genomic alterations and related tumor microenvironment is the biggest obstacle in development and identifying efficient cancer therapy^1^. To treat cancer, it is necessary to have many highly specific drugs and developed accurate diagnostics for targeted assignment of particular drugs or, more likely, combinations of drugs. Molecular profiling of tumors (TCGA ^2^) can provide the most complete data for construction of clinically relevant diagnostics, however, the molecular biomarkers for diagnostics can hardly be learned directly from even large clinical trials, because of the low success rate of the application of the specific drugs to tumors driven by diverse molecular alterations. This is especially true for emerging targeted immune therapies, which show fantastic results for a small percentage of patients and fails for the majority of cases ^3,4^. Despite multiple testing efforts and despite the fact that about 8% of primary tumors and 32% of metastatic castration-resistant tumors express PD-L1 ^5^, prostate cancer remains resistant to immunotherapy with PD-1/PD-L1 or CTLA-4 blockade (immune checkpoint blockade) ^6-10^. It has been proposed that combination therapy, including targeting potential compensatory mechanisms in immunotherapy-resistant prostate tumors, can potentially assist in the reactivation of anti-tumor immune response ^11,12^. Thus, it becomes clear that a success of treatment depends greatly on developed diagnostics for accurate prediction of a sensitivity of a given tumor to a particular therapy. However, it still remains unclear how to identify tumors, which will likely respond to checkpoint therapy, how to identify tumors, which will respond to available combination therapies, and what new combination therapies have to be developed to treat the remaining tumors?

To fill this clinical gap, we propose development of a hypothesis-driven diagnostic framework, which combines biological knowledge, analysis of large sets of cancer molecular profiles and prediction algorithms that can be applied to molecular data of a particular tumor. In this approach, molecular data of tumors linked to the results of clinical trials can be used to test hypotheses, and facilitating the identification of tumors sensitive to a particular targeted therapy. Currently, there is no accurate analytic protocol for determining which tumors will respond to checkpoint inhibition, and also for determining what combination therapy should be used to sensitize a tumor to treatment.

In this study, we relied on the assumption that the sensitivity of prostate cancer tumors to immune therapy can be deduced from expression levels of genes of the relevant molecular pathways. We assumed that computational analysis of large RNAseq datasets of primary prostate cancers of TCGA ^2^ and recently published metastatic set of MSKCC ^13^ can help narrow down a list of potential biomarkers that can be tested for prioritization of tumors for immune therapy. Specifically, in the development of the computational protocol for constructing molecular diagnostics for checkpoint immune therapy, we focused on the following questions: - How should tumors be ranked and optimally classified into potentially “sensitive” and “non-sensitive” groups? What are the relevant genes for assessing tumor sensitivity to a particular therapy? Should one use only a target gene or an entire molecular pathway? If a molecular pathway is chosen, what is the optimal scoring function for assessment of an activity of this pathway? And, finally, how can tumor classification be performed in practical setting, i.e. what are biomarkers and classification signatures that can be tested in practice? Answering these questions resulted in development the computational protocol for ranking tumors by sensitivity to checkpoint immune therapy; application of this protocol to molecular profiles of prostate cancer is the focus of this work.

## 2. Results

### IFN-γ/PD-L1 biological axis

Multiple studies showed that the immune responsiveness of tumor cells to immune checkpoint blockade is driven by interferon-gamma (IFN-γ) signaling ^14,15^. Interferon-gamma (IFN-γ) is a cytokine released by activated T cells into the tumor microenvironment (TME) in response to neoantigens presented on tumor cells ^11,16^. IFN-γ has both immunostimulatory and immunosuppressive functions that modulate the TME ^17^. Disruption of the balance between stimulatory and suppressive mechanisms of IFN-γ signaling can result in tumor immune evasion ^18^. Loss-of-function and genomic mutations in IFN-γ signaling genes were found associated with immune checkpoint blockade in clinical studies ^19-21^ and in genetic screens ^22-24^. Chronic IFN-γ signaling in tumor cells increased resistance to immune checkpoint blockade by multiple inhibitory pathways ^25^.

The IFN-γ signaling is driven by binding of IFN-γ from the microenvironment to tumor IFN-γ receptor subunits IFNGR1 and IFNGR2, inducing expressions of MHC-I ^26^ and PD-L1 (CD274) ^27^ (Fig. 1). Binding of IFN-γ to its receptor results in recruitment and activation of the Janus kinases, JAK1 and JAK2, and subsequent phosphorylation, dimerization, and activation of a transcription factor known as signal transducer and activator of transcription, STAT1. STAT1 homodimers then translocate to the nucleus, where they bind to specific promoter elements and modulate transcription of IFN-γ-regulated genes ^15,16,28^.

**Figure 1.**
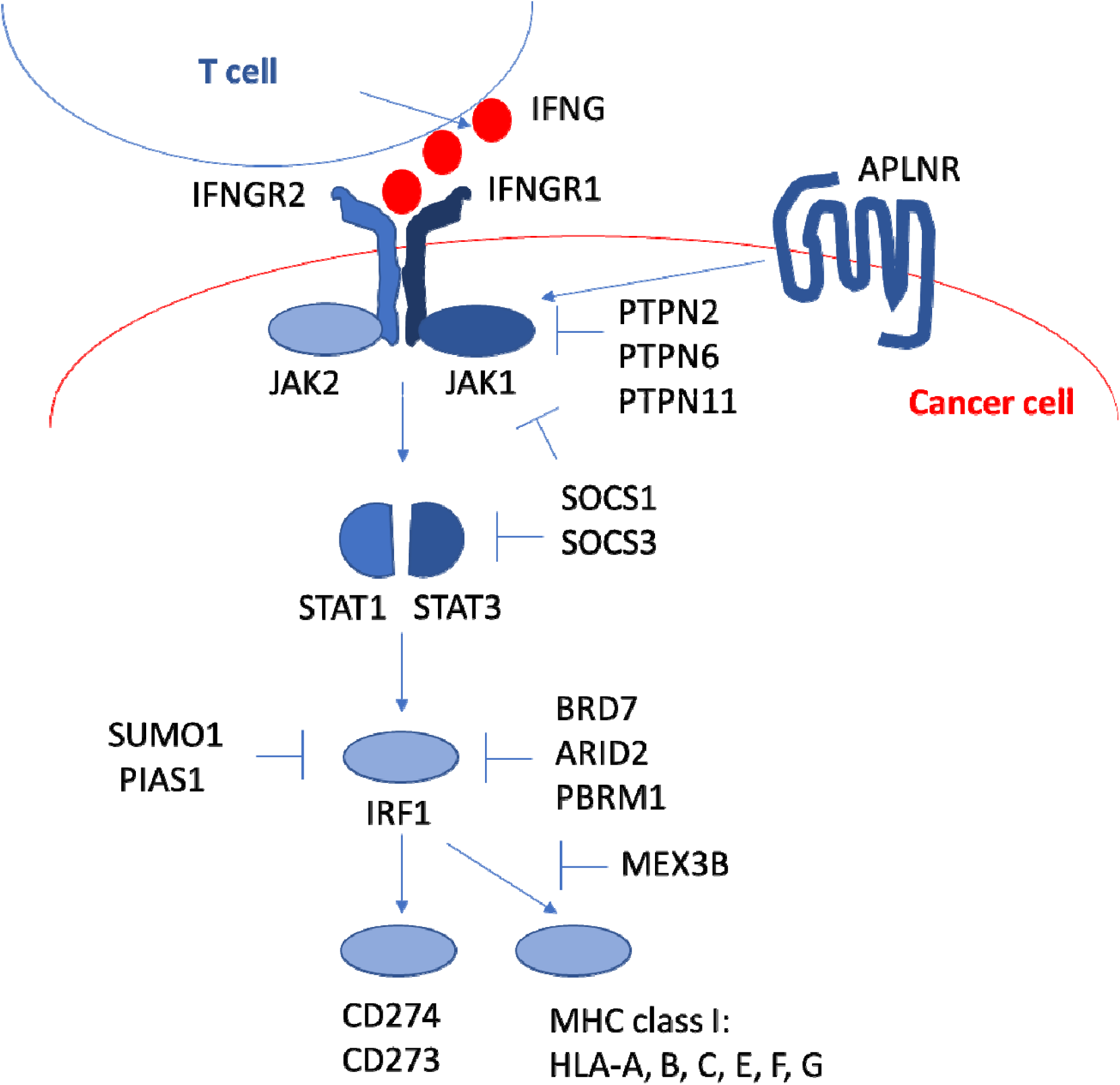
Schematic of IFN-γ/PD-L1 signaling axis. A reactive T cell recognizes a cancer cell antigens via MHC class molecules and releases IFN-γ (IFNG) that binds by the IFN-γ receptors (IFNGR1, INFGR2) on the cancer cell resulting in activation of the JAK-STAT pathway (JAK1, JAK2, STAT1, STAT3). This activates the transcription factor IRF1, which then activates the transcription of CD274 (PD-L1 is a product of CD274). Expression of PD-L1/2 on the surface of the cancer cell negatively regulates the anti-cancer T cell response ^14,15^. IFN-γ signaling is positively regulated by the above genes and by receptor apelin (APLNR) and negatively regulated by non-receptor tyrosine-protein phosphatases PTPN2, PTPN6, PTPN11 ^22-24^ and by suppressor of cytokine signaling (SOCS) proteins ^70^. Transcription of IFN-γ target genes is negatively regulated by the chromatin remodeling complex PBAF (BRD7, ARID2, PBRM1) ^71^ and SUMO, PIAS1 ^72^. IFN-γ dependent transcriptional regulation of MHC class I (HLA-A) is inhibited by MEX3B ^73^.

Thus, based on the key role of IFN-γ in modulation of immune and inflammatory responses and tumor immunosurveillance, we chose “IFN-γ/PD-L1 axis” as the primary pathway controlling tumor sensitivity to PD-L1/2 therapy - (Fig.1) We settled with this signaling pathway because it has all necessary components – a cytokine mediator, receptors, signal transduction, transcription factors and final effectors (PD-L1/2, MHC-I), and all those genes are expressed in cancer cells. We hypothesized that non-responder tumors are either tumors with downregulation of IFN-γ axis genes, which make them invisible to immune cells, or tumors with upregulated IFN-γ axis genes and another specifically activated mechanism of immune evasion preventing cancer cell death. Then, ranking tumors by activation of IFN-γ axis enabled us to classify tumors by sensitivity to checkpoint inhibition.

### IFN-γ axes and methods for inferring pathway activity

Comparison of axes of a single CD274, positive regulators of IFN-γ/PD-L1 pathway and combination of positive and negative regulators of IFN-γ/PD-L1 pathway are presented in Table 1.

**Table 1.** Statistics of Pearson correlation coefficients computed between gene expression levels of prostate cancer tumors and scoring functions representing activation of IFN-γ/PD-L1 axis^$^.

| Correlation threshold /<br>Scoring functions<sup>\$\$</sup> | primary tumors (TCGA) | | | | | | | | | metastatic tumors (MSKCC) | | | | | | | | |
| --- | --- | --- | --- | --- | --- | --- | --- | --- | --- | --- | --- | --- | --- | --- | --- | --- | --- | --- |
|  | 0.1 | 0.2 | 0.3 | 0.4 | 0.5 | 0.6 | 0.7 | 0.8 |  | 0.1 | 0.2 | 0.3 | 0 | 0.5 | 0.6 | 0.7 | 0.8 |  |
| CD274 | 10337 | 5357 | 1819 | 732 | 401 | 170 | **36** | 0 |  | 475 | 80 | 18 | 9 | 5 | 4 | 2 | 0 |  |
| IPAS(IFNG<sub>P</sub>) | 11461 | **6409** | **2912** | 1042 | **511** | **189** | 17 | 0 |  | 8243 | 4748 | 2156 | 731 | **292** | **107** | **15** | 0 |  |
| ssGSEA(IFNG<sub>P</sub>) | **11543** | 6406 | 2876 | **1070** | 466 | 119 | 8 | 0 |  | **8446** | **4822** | **2256** | **748** | 226 | 61 | 5 | 0 |  |
| <IFNG<sub>P</sub>> | 8228 | 2579 | 867 | 398 | 115 | 24 | 11 | **6** |  | 7116 | 2962 | 938 | 330 | 116 | 20 | 3 | 0 |  |
| IPAS(IFNG<sub>P</sub>) - IPAS(IFNG<sub>N</sub>) | 9620 | 4017 | 1502 | **751** | **383** | **82** | **14** | **1** |  | **9116** | **4383** | **1755** | **520** | **158** | **24** | 0 | 0 |  |
| ssGSEA(IFNG<sub>P</sub>) - ssGSEA(IFNG<sub>N</sub>) | **10735** | **5302** | **1614** | 193 | 19 | 2 | 0 | 0 |  | 8221 | 3798 | 1497 | 363 | 45 | 0 | 0 | 0 |  |
| <IFNG<sub>P</sub>> - <IFNG<sub>N</sub>> | 8516 | 2722 | 818 | 345 | 92 | 21 | 10 | 6 |  | 7059 | 2895 | 870 | 299 | 102 | 17 | 3 | 0 |  |

<sup>\$</sup>Numbers of genes with the absolute value of Pearson correlation coefficient computed between gene's expression levels of TCGA and MSKSS sets of prostate tumors and values of the pathway scoring functions above or equal to given correlation thresholds (0.1, 0.2,...0.8); <sup>\$\$</sup>Scoring functions used in computing correlations: CD274 stands for expression levels of the gene CD274; IPAS(IFNG<sub>P</sub>) and ssGSEA(IFNG<sub>P</sub>) stand, respectively, for IPAS and ssGSEA scoring functions computed for genes, which positively regulate expression of CD274 in IFN-γ pathway of Fig.1; <IFNG<sub>P</sub>> represent average expression levels of positive regulators of IFN-γ pathway of Fig.1; the combined activity of IFN-γ pathway, which include both positive and negative regulators, was represented by the differences of IPAS and ssGSEA scoring functions computed, respectively, for sets of positive (IFNG<sub>P</sub>) and negative (IFNG<sub>N</sub>) regulator genes and also by the difference of average gene expression levels of positive, <IFNG<sub>P</sub>> , and negative, <IFNG<sub>N</sub>>, regulators; ; the maximum numbers of correlated genes obtained for each of thresholds are shown in bold.

We used three methods for inferring pathway activity across tumor samples: averaged expression of pathways genes and two enrichment scores ssGSEA and IPAS (Supplement 1). A combination activity of a pathway with positive and negative components was assessed as a difference between independently assessed activities of positive and negative components. For better comparisons, we put together correlation distributions computed for both primary and metastatic tumors. As a reference axis, we took the ordering of tumors by expression level of CD274 genes. There is a drastic difference between primary and metastatic tumors revealed by a number of genes correlated with expression level of CD274. With an increase of correlation threshold, only a few highly correlated genes remain in metastatic cancers as compared to hundreds in primary tumors. For positive regulators, the enrichment scores have significantly more correlations as compared to CD274 axis and average expression axis for both primary and metastatic tumors. Surprisingly, CD274 axis produced more correlated genes as compared to the axis formed by averaging expression levels. The IPAS axis produced more highly correlated genes as compared to ssGSEA axis for both primary and metastatic tumors. The similar trend is seen in combination of scoring functions computed for positive and negative components of IFN-γ/PD-L1 pathway. However, the decrease of highly correlated genes is stronger for ssGSEA score as compared to the averaging score.

Thus, ranking tumors along IFN-γ/PD-L1 signaling axis using the pathway enrichment scores revealed more correlated genes and, in that way, more biological differences between tumors as compares to ranking tumors along the axis of CD274 expression level or the axis formed by averaging expression levels. Thus, the pathway scoring functions can be more informative for diagnostics as compared to the use of single biomarkers. The IPAS score revealed more high correlated genes as compared to ssGSEA score. Therefore, in further analysis – revealing potential immune subtypes, characteristic genes and pathways (Fig.2) we used IPAS scores for ranking tumors along IFN-γ/PD-L1 signaling axis.

### Determining immune subtypes

We proposed to determine disease subtypes by determining a point of the maximal difference between two tumor sets aligned along IFN-γ/PD-L1 signaling axis (Methods, Fig.6, Supplement 2). We assumed here that the true biologically different disease subtypes have to have the maximal difference between gene expression distributions. However, position of such a point of the maximal difference may depend on a number of genes used in computation of the combined score (Supplement 2). Therefore, following the protocol of Fig.6, we examined different gene sets filtered by the absolute value of correlation with the IFN-γ/PD-L1 axis score (Fig.2). For primary prostate tumors, when all genes were taken into account (correlation threshold 0); we found a distinct minimum, which separated the most significantly IFN-γ-depleted tumors from the rest (Fig.2A). With increase of the correlation threshold, the combined scoring function became rather shallow, while preserving position of the minimum (Fig.2A). Thus, for primary tumors, we considered two separation into IFN-γ-depleted (“cold”) and IFN-γ enriched (“hot”) subtypes. The first separation defines “truly” IFN-γ-depleted tumors; the second separation defines IFN-γ-enriched tumors. For metastatic prostate tumors, the scoring functions remained shallow and pointed to the same position of the minimum for all correlation thresholds up to 0.5 (Fig.2B). At correlation threshold of 0.6, position of the global minimum had shifted to right along IFN-γ axis and a new local minimum emerged in IFN-γ-enriched region. Because a number of correlated genes at threshold of 0.6 was small, we decided to choose for current analysis the consistent position of the minima obtained for correlation threshold of 0.0-0.5 as in Fig.2B.

**Figure 2.**
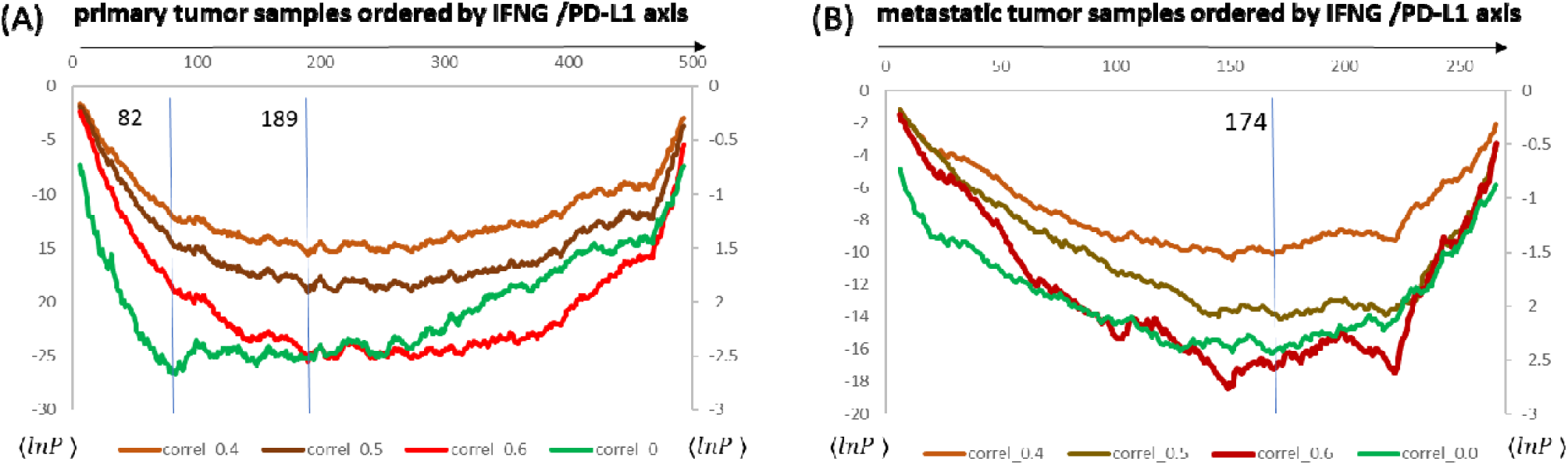
Determining position of the maximal difference between IFN-γ-depleted and IFN-γ-enriched subtypes for primary and metastati**c** tumor sets. We identify two IFN-γ-depleted subtypes of 82 and 189 tumors of total 497 primary tumors **(A)** and a subtype of 174 tumors of 270 of total metastatic tumors **(B)**. The position of the maximal difference between tumors ranked by increase of inferred activity of IFN-γ/PD-L1 signaling is determined by the minimum of the combined scoring function (Supplement 2). We compare two axes: the positive IPAS (IFNG_P_) and combined IPAS (IFNG_P_) – IPAS (IFNG_N_). The combined axis makes deeper separation of tumors (lower value of the combined scoring function). The genes of IFN-γ/PD-L1 pathway are omitted in computations to minimize the bias from the axis genes. The right vertical axis corresponds to green plots when all genes are taken into account (correlation threshold equal to zero); the depth of the minimum increases for sets of higher correlated genes **(A, B)**. The vertical blue lines show positions of the determined minima **(A, B)**.

### Identification of key genes, pathways and cell type signatures for primary prostate tumors

Prostate cancer is considered poorly immunogenic and characterized by low tumor infiltration with cytotoxic T-cells ^29^.To gain insight into the mechanisms of low immunogenicity and immune evasion in prostate cancer, we analyzed genes and pathways in primary tumors with the highest increase of inferred activation of the IFN-γ/PD-L1 axis. We compared two groups ranked high or low along the IFN-γ/PD-L1 axis, which we labeled “IFN-γ-depleted” or “IFN-γ-enriched” tumor subtypes, respectively. From this analysis, we observed that IFN-γ-enriched primary tumors are inflamed and enriched with genes and pathway involved in multiple immune evasion mechanisms.

In the IFN-γ-enriched subtype we obtained 426 genes and 714 pathways including prostate specific pathways: prostate cancer pathways (p = 9.1E-12), miRNA regulation of prostate cancer signaling pathways (p = 5.6E-06) (cutoff, 2-fold change (FC) with p < 0.00001). Among the top 200 genes the highest increase was identified in genes involved in signal transduction (30 genes), immune system (28 genes) and in inflammatory response (24 genes). In particular, we found genes mediating G-protein coupled receptor, cytokine- and chemokine-mediated signaling pathways (27 genes) and neutrophil degranulation (16 genes). Among the top 50 pathways the highest increase was detected in the immune system pathway, neutrophil degranulation, cytokine and GPCR signaling. The identified pathways were consistent with the functions of the top 200 upregulated genes demonstrating the inflammatory environment of the IFN-γ-enriched subtype.

We also identified upregulated immune checkpoints in IFN-γ-enriched tumors, known to reduce inflammation under normal physiological conditions and that are hijacked during tumor immune evasion ^30^. We found increase in canonical immune checkpoints: CTLA4 inhibitory signaling pathway (p= 2.3E-10), CTLA4 gene (2.4FC, p = 0); PD-L1 expression and PD-1 checkpoint pathway in cancer (p = 0), genes CD274 (PD-L1) (2FC, p = 0) and PDCD1LG2 (PD-L2) (2FC, p = 0). Among alternative immune checkpoints, the most upregulated were TIGIT and BTLA (both 3FC, p = 0) and VISTA (VSIR) gene (2FC, p = 0); LAG3 was 2FC, p = 4.4E-13.

**Figure 3.**
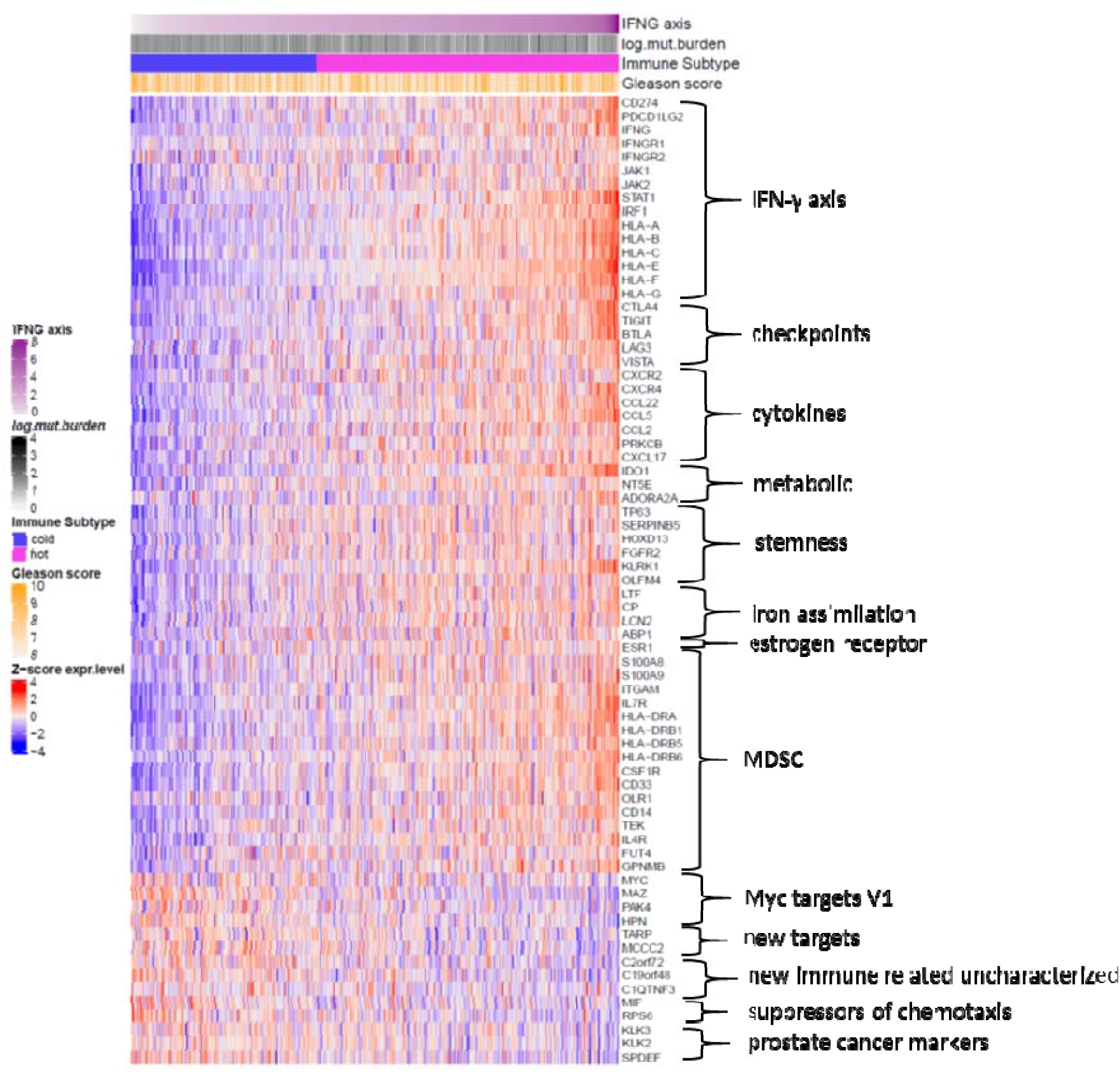
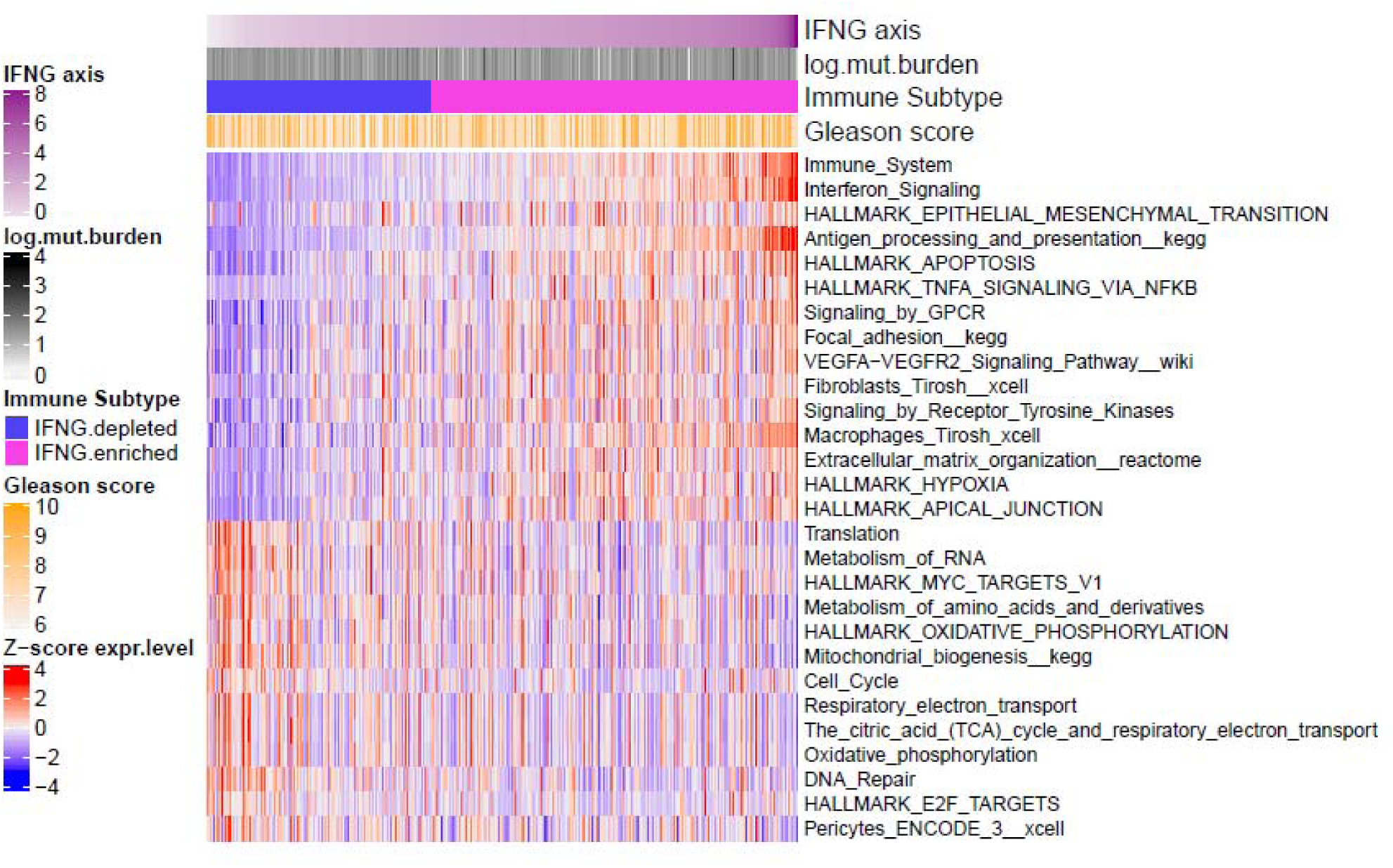
(A) Heatmap of major genes with known or proposed role in immune evasion determined as differentially expressed between IFN-γ-enriched and IFN-γ-depleted subtypes of primary prostate cancers. (B) Heatmap of major pathways determined as differentially activated between IFN-γ-enriched and IFN-γ-depleted subtypes of primary prostate cancers.

**Figure 4.**
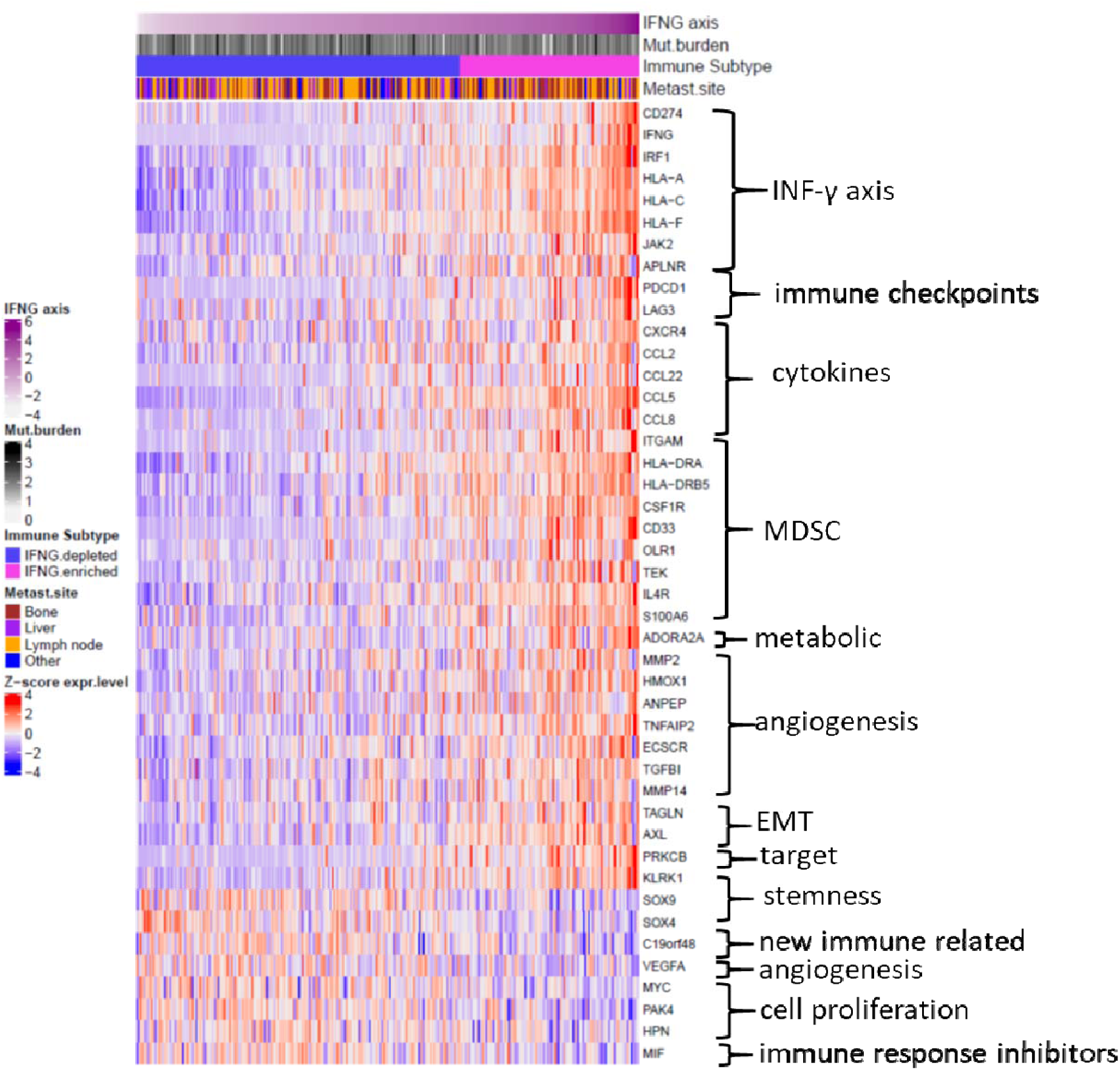
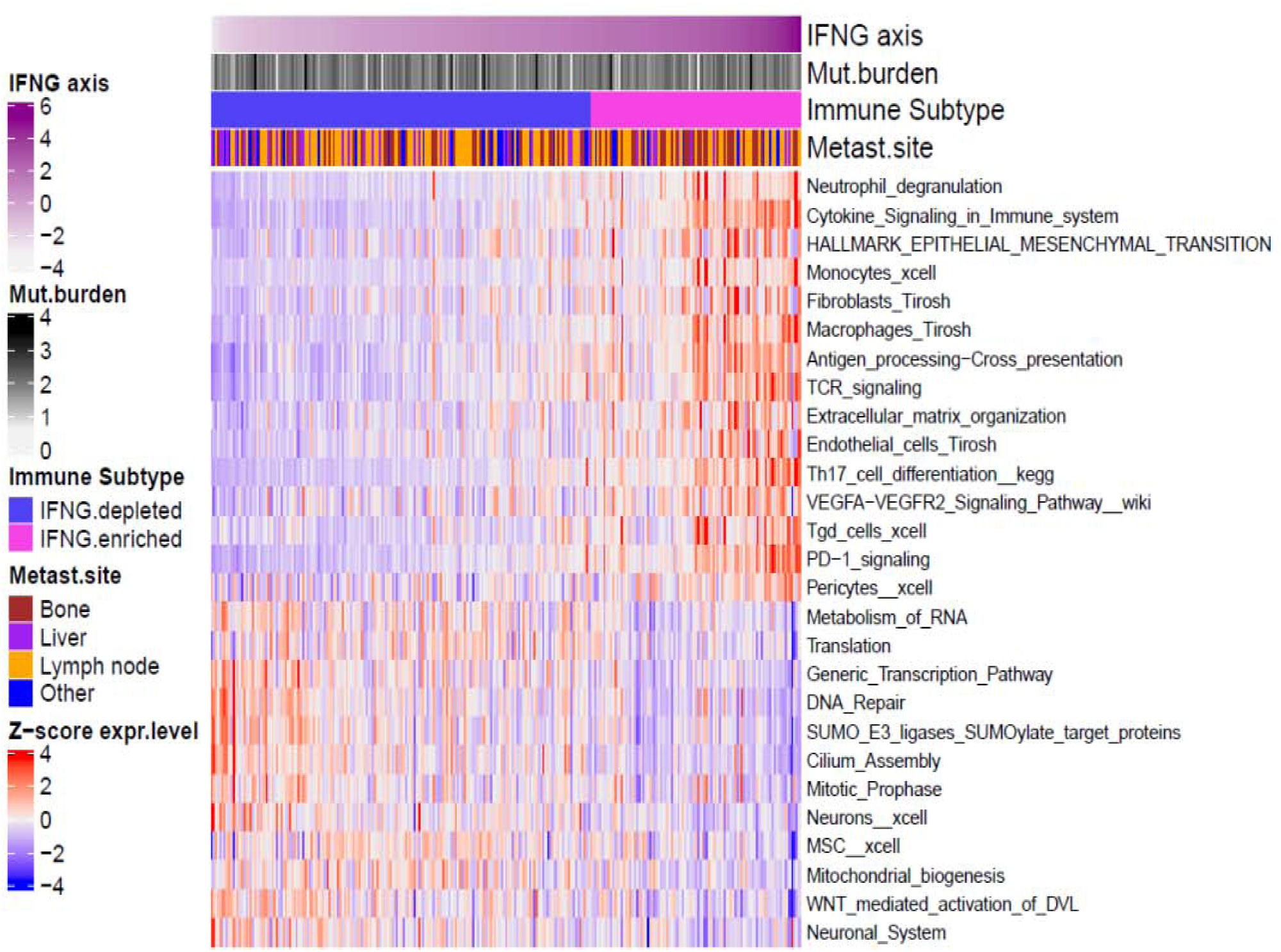
(A) Heatmap of major genes with known or proposed role in immune evasion determined as differentially expressed between IFN-γ-enriched and IFN-γ-depleted subtypes of metastatic prostate cancers. (B) Heatmap of major pathways determined as differentially activated between IFN-γ-enriched and IFN-γ-depleted subtypes of metastatic prostate cancers.

**Figure 5.**
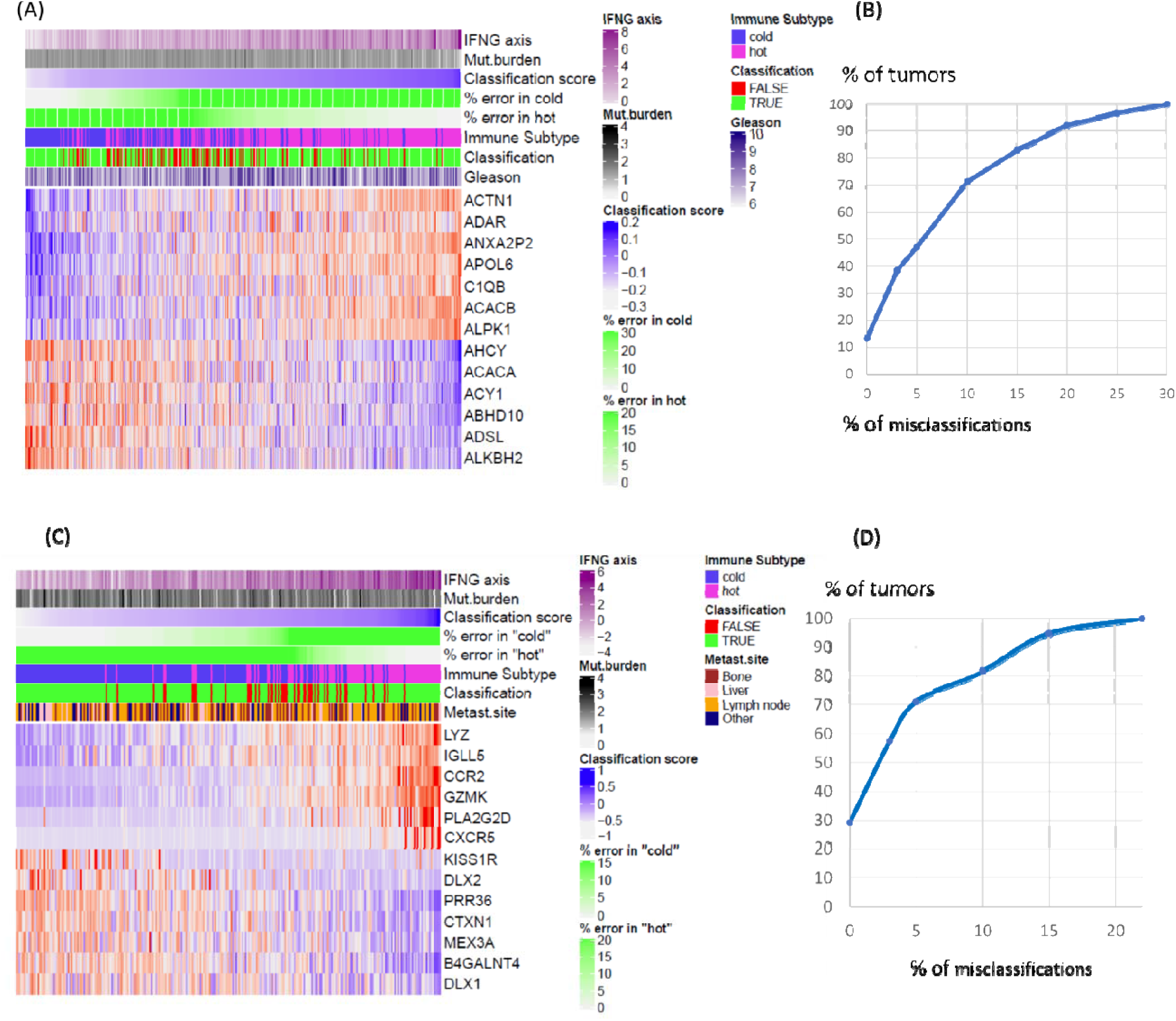
Biomarker genes proposed for classification of primary and metastatic prostate tumors. Heatmaps of expression levels of proposed biomarker genes for classification of primary and metastatic prostate tumors **(A, C)** and related classification summary plots **(B, D)**. The classification scores were computed using IPAS method (Supplement 2) applied to “positive” and “negative” biomarker genes. “Positive” and “negative” biomarkers were defined based on, respectfully, significant upregulation or downregulation of expression levels in IFN-γ enriched immune subtype as compared to “IFN-γ-depleted” immune subtype. Thus, given only relative positions of “positive” and “negative” genes in a set of 14 (primary) or 11 (metastatic) biomarker genes, one can classify tumors based on a value of the IPAS score. Panels **(C)** and present percentages of tumors which were classified within a given misclassification threshold; in particular, ~50% of primary tumors and ~70% of metastatic tumors were classified in immune IFN-γ-enriched or IFN-γ-depleted subtypes with percentage of misclassifications less than 5%. As accuracy of classification depends on a value the classification score, percentages of misclassifications were computed separately for each of the immune subtypes as function of the classification score. Selecting genes for the panel, we used the following criteria: genes upregulated in IFN-γ-enriched or IFN-γ-depleted subtypes were sorted by P values (<0.0001) and then by fold changes (>2), and, finally, we excluded genes with very low or very high average expression level.

Resistance to the immune checkpoint blockade is associated with the tumor microenvironment enriched in immune suppressive cells and cytokines ^31^. We identified gene signatures of multiple cell types in the IFN-γ-enriched subtype with the highest upregulation in gene signature of fibroblasts (p = 1.84E-11, Fibroblasts Tirosh xcell complete signatures) and macrophages (p = 0, Macrophages Tirosh xcell complete signatures) with both populations of M1 (p = 0) and M2 (p = 7.19E-13). We found upregulation of immunosuppressive cytokines that recruit regulatory T cells (Tregs) and myeloid-derived suppressor cells (MDSC) (CXCR2 2.4FC, p = 8.88E-16; CXCR4, 2FC, p = 0; CCL22, 2.4FC, p = 0, CCL5, 2.5FC, p = 0) that was consistent with increase in regulatory T cell population (Tregs Charoentong xcell complete signatures, p = 0). We observed the prevalence of immunosuppressive cells over cytotoxic lymphocytes (CD8+ Tem BLUEPRINT 3 xcell, p = 0; CD8+ Tem Charoentong xcell complete signatures, p = 0; CD4+ Tem NOVERSHTERN 1 xcell, p = 0; CD4+ Tcm Charoentong xcell complete signatures, p = 0; NK cells FANTOM 1 xcell, p = 0; NK cells Rooney xcell complete signatures, p = 8.4E-12; Natural killer cell mediated cytotoxicity pathway, p =0).

Among known immunosuppressive pathways associated with resistance to immunotherapy through recruitment of immature dendritic cells, MDSC and Treg cells, we detected upregulation of VEGF signaling pathway ^30^ VEGFA-VEGFR2 Signaling Pathway, p = 2.57E-13, PRKCB (2FC, p = 3.3E-16), CCL2 (2.4FC, p = 0) and CXCL17 (4.8FC, p = 0), TGF-beta signaling pathway (p = 9.9E-09) and IL-10 anti-inflammatory signaling pathway (p = 0), IL10 (2FC, p = 4.4E-16). Significant upregulation of CXCL17 positioned this gene among the top three upregulated genes along the IFN-γ/PD-L1 axis. CXCL17 regulates T cell homeostasis and it is expressed under inflammatory conditions functioning as a chemoattractant for macrophages, dendritic cells and is capable of attenuating inflammation through the recruitment of myeloid derived suppressor cells ^32,33^.

Tumor cell-intrinsic signaling pathways also shape the tumor immune landscape ^15^. We identified upregulation of IFN-γ responsive genes IDO1 (3.3FC, p = 0) and TDO2 (2.8FC, p = 3.1E-8), responsible for activation of MDSCs and suppression of T cells. As tumor inflammation causes hypoxia we detected significant upregulation in hypoxia (Hypoxia, p = 0) positioned among the top 50 activated pathways. One of the consequences of hypoxia is suppression of T cell function by upredulation of adenosine-mediated inhibitory pathway of immune evasion and consistent with this hypothesis, we identified upregulation of genes NT5E, ADORA2A.

Interestingly, upregulation of genes involved in development of the prostate gland and genitalia TP63, HOXD13, FGFR2 and Developmental Biology pathway in the IFN-γ-enriched subtype indicates on the involvement of tumor dedifferentiation and possible stemness mechanisms of immune evasion ^15^.

Alteration in iron availability and iron metabolism can be immunosuppressive leading to tumor immune evasion ^34^. Among the top 15 genes of IFN-γ/PD-L1 axis, four of them were genes regulating iron transport: LTF, 7FC, p = 0; CP, 4.2FC, p = 0; LCN2, 3.8FC, p = 1.1E-16; ABP1 (AOC1), 3.7FC, p = 3.5E-11. Consistent with activation of the iron homeostasis pathways (Mtb iron assimilation by chelation (p = 0); iron uptake and transport (p = 0), LTF danger signal response pathway (p = 0), observed upregulation in the IFN-γ-enriched subtype indicates the iron depletion in the tumor microenvironment. Iron deficiency impairs immune response ^35^ including anti-tumor immunity ^36^. Similar to the oxygen depletion (hypoxia), deficiency in iron activates HIF1-alpha and its responsive genes required for cell survival under stress ^37^. As iron is needed for IFN-γ-induced tumor ferroptosis ^38^ it was proposed to target this pathway in combination with immune checkpoint blockade treatment.

These data confirm presence of established immune suppressive cell-mediated mechanisms of resistance to immunotherapy in the IFN-γ-enriched subtype and reveal genes with anti-inflammatory, immunoregulative and protective activities such as CXCL17 ^33^, LTF ^39^ and CP ^36,37^ as promising anti-tumor immune therapy targets in this subtype.

On the opposite side of the IFN-γ/PD-L1 axis, in the IFN-γ-depleted subtype we obtained 200 genes and 446 pathway (cutoff, 1.3-fold change (FC) with p < 0.00001). Among 200 genes, the highest increase (2.2 - 1.3FC) was in genes involved in translation (13 genes), rRNA processing (14 genes), translation initiation (13 genes), transcription (13 genes), oxidation-reduction process (8 genes) and signal transduction (8 genes). Pathway with the highest increase (18K - 6FC) were translation, RNA metabolism and cell cycle. We found a number of pathways driving translation with the initiation step in translation at the top of the list (Cap-dependent translation initiation pathway, p = 2.7E-06). The critical role of transcription initiation in prostate tumorigenesis is recognized and currently under investigation ^40-44^. We identified multiple oncogenic factors that were translationally regulated in the IFN-γ-depleted subtype including cell cycle control and anti-apoptotic molecules (MYC targets V1 pathway (p = 7.4E-06), Cell Cycle pathway (p = 9.6E-07), genes MYC, NKX3.1, SOX9, TBX3, CDK19, PAK4, HPN).

Increased tumor cell oxidative metabolism is associated with repressed T cell function during T cell tumor infiltration and poor survival after anti-PD-1 immunotherapy in melanoma ^45^. Consistent with the hypothesis that oxidative axis represents an important barrier to effective immunotherapy we found significant upregulation of oxidative phosphorylation in the IFN-γ-depleted subtype: oxidative phosphorylation pathway (p = 8.8E-04), mitochondrial biogenesis (p = 1.5E-11), respiratory electron transport (p = 2.6E-05).

Due to the narrow fold change range (2.2-1.5FC), arrangement by the highest average expression revealed the TARP gene (1.5FC, p = 9.0E-09). As a highly expressed protein in primary and metastatic prostate tumors, TARP was used as a tumor antigen to develop a successful vaccine that increased patient survival ^46^. The other top list gene was MCCC2 (1.5FC, 5.0E-08), methylcrotonoyl-CoA carboxylase 2, which is involved in tumor formation ^47,48^.

We found a number of uncharacterized proteins with prognostic and therapeutic potential (genes: LOC100130872, C2orf72, C19orf48). In particular, C19orf48 encodes a multi-drug resistance protein overexpressed in androgen receptor-dependent manner in prostate cancer ^49^. C19orf48, a minor histocompatibility antigen recognized by cytotoxic T cells was demonstrated to have an antitumor effects in kidney cancer patients ^50^.

We identified significant enrichment in the gene signature for pericytes (p = 8.1E-05). Due to expression of PD-L1/2 and immunosuppressive properties (NO, PGE2, and TGF-β), pericytes drive immune evasion in gliomas ^51^. The angiogenic role of pericytes in prostate tumor microenvironment ^52,53^ in the IFN-γ-depleted subtype deserves further investigation.

Among top 200 genes with highest fold change we also identified upregulated genes involved in negative regulation of immune response (C1QTNF3), in activation-induced cell death of T cells (RPS6), negative regulation of macrophage chemotaxis (MIF), VEGF and angiogenesis (MAPK12, ADM2), that contribute to the phenotype of the IFN-γ-depleted tumors.

### Characteristic genes and pathways obtained for metastatic tumors

Tumor-induced systemic inflammation has been associated with metastasis in multiple cancers including prostate cancer ^54^. To understand how the status of IFN-γ affects immune evasion mechanisms of metastatic prostate cancer, we compared the transcriptomes of the two groups – metastatic tumors of the IFN-γ-enriched subtype with metastatic tumors of the IFN-γ-depleted subtype.

In the IFN-γ-enriched subtype in metastatic tumors we obtained 203 genes and 776 pathway (cutoff, 2-FC with p < 0.00001). The transcriptome of the top 200 genes of the IFN-γ-enriched subtype was dominated by genes regulating neutrophil degranulation (37 genes), which is consistent with an increased pool of neutrophils due to the systemic inflammation ^55^ and with the promoting role of neutrophils in metastatic spread ^56^. Among the top 200 genes were also genes promoting tumor cell invasion and metastasis formation including cell adhesion (16 genes), proteolysis (15 genes), extracellular matrix organization and degradation (12 genes), leukocyte migration (11 genes), negative regulation of apoptosis (11 genes), cell-cell signaling (10 genes), cell migration (10 genes) and angiogenesis (10 genes). Among the top upregulated 50 pathways, we found significant increase in the fold change of the neutrophil degranulation (p = 0), cytokine signaling (p = 0), epithelial-to-mesenchymal transition pathway (p = 6.3E-11), antigen processing-cross presentation (p = 0), TCR signaling (p = 0), extracellular matrix organization (p = 7.3E-15).

Among established immune evasion mechanisms contributing to metastatic disease, we identified significant increase in selected immune checkpoints including PD-1 (PDCD1, 3.6FC, p= 1.9E-15), PD-L1 expression and PD-1 checkpoint pathway in cancer (p = 0), VISTA (VSIR) gene (3.2FC, p = 0) and in CD274 (PD-L1) (2.3FC, p = 2E-13). We observed upregulation of epithelial-to-mesenchymal transition genes TAGLN (2.1FC, p = 5.5E-8) and AXL (2.3FC, p =0) involved in resistance to the immune checkpoint blockade^57^.

The features of TME of the metastatic IFN-γ-enriched subtype included enrichment in fibroblast gene signature (Fibroblasts Tirosh xcell complete signatures, p = 2.6E-11) and increase in macrophages (Macrophages Tirosh xcell complete signatures, p = 0) and monocytes (Monocytes FANTOM 2 xcell, p = 0) and their chemotactic genes CCL8 (3FC, p = 3.8E-13) and inhibitory cytokines (CXCR2, 3.3FC, p = 2.0E-08), CCL5 (5FC, p = 0). We also identified signatures of endothelial cells (Endothelial cells Tirosh xcell complete signatures, p = 1.6E-14), Th17 cells (Th17 cell differentiation pathway, p = 0), Tγδ cells (Tgd cells HPCA 3 xcell, p = 2.5E-11) and pericytes (p = 5.9E-06).

Upregulation of signaling by VEGF (p = 3.9E-13), VEGFA-VEGFR2 pathway (p = 7.9E-13), VEGFA gene (1.7FC, p =3.3E-06) and angiogenesis genes (TGFBI, CCL2, MMP14, MMP2, ANPEP, ECSCR, APLNR, HMOX1 and TNFAIP2, all 2-3FC with p < E-07) indicate immunosuppression mechanism by VEGF in addition to promotion of tumor angiogenesis ^30^.

In summary, these data reveal the active role of the immunosuppressive tumor microenvironment with the major role of the immunosuppressive myeloid cells in modulation of immune response in metastatic IFN-γ-enriched subtype of prostate tumors.

IFN-γ-depleted subtype in metastatic tumors we obtained 215 genes and 303 pathway (cutoff, 1.5-FC with p < 0.001). The top 200 upregulated genes were involved in transcription (39 genes), cell proliferation (27 genes), signal transduction (16 genes), oxidation-reduction process (12 genes), cell adhesion (12 genes) and hypoxia (8 genes). Among the top 50 pathways we identified metabolism of RNA (p = 3.8E-06), translation (p = 3.3E-05), transcription (p = 3.4E-11), cell cycle (p = 1.1E-03), mitochondrial biogenesis (p = 1.8E-08), DNA repair (p = 1.8E-08), ubiquitination (p = 6.7E-10) and cilia assembly (p = 2.6E-14).

We identified upregulation of the Wnt signaling pathway (WNT mediated activation of DVL pathway p = 1.8E-07), genes DVL2 (1.3FC, p = 1.2E-05), AXIN1 (1.3FC, p = 8.5E-05)), and its positive regulator SOX4 (1.7FC, p = 2.9E-08) and its mediator PAK4 (1.7FC, p = 7E-08) confirming the role of increased Wnt signaling in developing resistance to immune checkpoint blockade ^15^.

Upregulation of SOX9 gene (1.7FC, p = 6.1E-06) in the metastatic IFN-γ-depleted subtype is consistent with its role in promoting metastasis in prostate cancer ^58^. SOX9 is an important regulator of prostate gland development and its overexpression in advanced lung adenocarcinoma metastasis confers resistance to natural killer-mediated cell killing ^59^ pointing toward SOX9-mediated evasion of immune response in the metastatic IFN-γ-depleted subtype. These data demonstrate the presence of immune evasion mechanisms in the metastatic IFN-γ-depleted subtype that are mediated by upregulation of Wnt signaling and stem-like program including SOX9.

We identified pericytes cell type signature (Pericytes ENCODE 3 xcell p = 0.5E-04)and mesenchymal stem cell signature (MSC HPCA 3 xcell, p = 2.4E-04). Identified neuronal signature (Neurons ENCODE 2 xcell, p = 8.0E-07) in the metastatic IFN-γ-depleted subtype indicates presence of perineural invasion^60^, an indicator of aggressive prostate cancer^61,62^.

### Classification of prostate tumors into IFN-γ-enriched and IFN-γ-depleted subtypes

To assign immune therapy correctly and, to assign, when necessary, additional specific drugs to neutralize immune evasion, one needs accurate diagnostics. The results presented in previous sections showed significant difference between IFN-γ-enriched and IFN-γ-depleted tumors both in primary and metastatic tumors. Therefore, it is not difficult to propose a set of biomarker genes for accurate classification of tumors by immune subtype or along IFN-γ/PD-L1 axis. However, the sequencing platforms and computational pipeline used to assess gene expression levels are subject to batch effect, while correction of batch effect needs large data sets that is problematic in practice. Therefore, typically, one cannot used directly expression levels of biomarker genes derived in one study to classification of transcriptional profiles produced in another study. We propose a solution to this problem by transition from profiles of gene expression levels to profiles of pathway activity scores. The multiple classification tests run with MSM ^63^ algorithm showed that ssGSEA and IPAS scoring methods, which assess collective up/down-regulation of pathway genes based on ranking genes in each of tumors, can be used in classification individual tumors given whole-genome transcriptional profiles (Supplement 3). However, getting whole-genome transcriptional profile for each of tumors, is not the best practical solution. - Can accurate diagnostics be developed with a smaller number of genes and with no complications caused by batch effect? – To this end, we propose biomarker panels composed of two gene sets, which are inversely regulated in IFN-γ-enriched and in IFN-γ-depleted subtypes, i.e. one “positive” gene set is upregulated in IFN-γ enriched and downregulated in IFN-γ-depleted subtype, another “negative” set is up-regulated in IFN-γ-depleted and down regulated in IFN-γ-enriched subtype. Thus, genes of such a panel can be treated as a pathway composed of positive and negative branches. Using the IPAS scoring method, we computed two scores, an activation score for a “positive set”, and a suppression score for a negative set, and used these scores to feed in our classification algorithm ^63^. Assessing classification accuracies, we simulated a practical situation, when based on a scoring value, one needs to classify a particular sample. In particular, we assessed accumulation of misclassification error independently for IFN-γ-depleted and IFN-γ enriched subtypes as functions of the classification score. Thus, for primary prostate tumors ~50% of tumors were classified correctly in IFN-γ-enriched and IFN-γ-depleted subtypes with error less than 5%; ~70% of tumors were classified correctly with error less than 10%;for metastatic tumors, ~70% and 80% of tumors were classified correctly with errors less than 5% and 10%, respectively. Taking into account that no special optimization was done in selection of biomarkers, we believe that the proposed approach has a significant potential for development accurate diagnostic panels with a small number of biomarker genes.

## 3. Methods

The schema of the computational protocol used in this work is presented in Fig.6. We proposed to order tumors along a “biological axis”, which can be presented by an expression level of a target gene (in our case, CD274) or by inferred activity of a relevant molecular pathway (a pathway score) or by experimentally derived sensitivities to drug treatment (IC50, EC50). Then, we (i) rank different biological axes by a number of correlated genes; (ii) select a subset of correlated genes for (iii) determining two most distinct sets of tumors (potential disease subtypes) separated by a position on a biological axis. Two-class separation of tumors make possible to determine specific genes and pathways – potential biomarkers - to better characterization biological differences and determine biomarkers for classification of tumors along the axis (Supplement 2).

**Figure 6.**
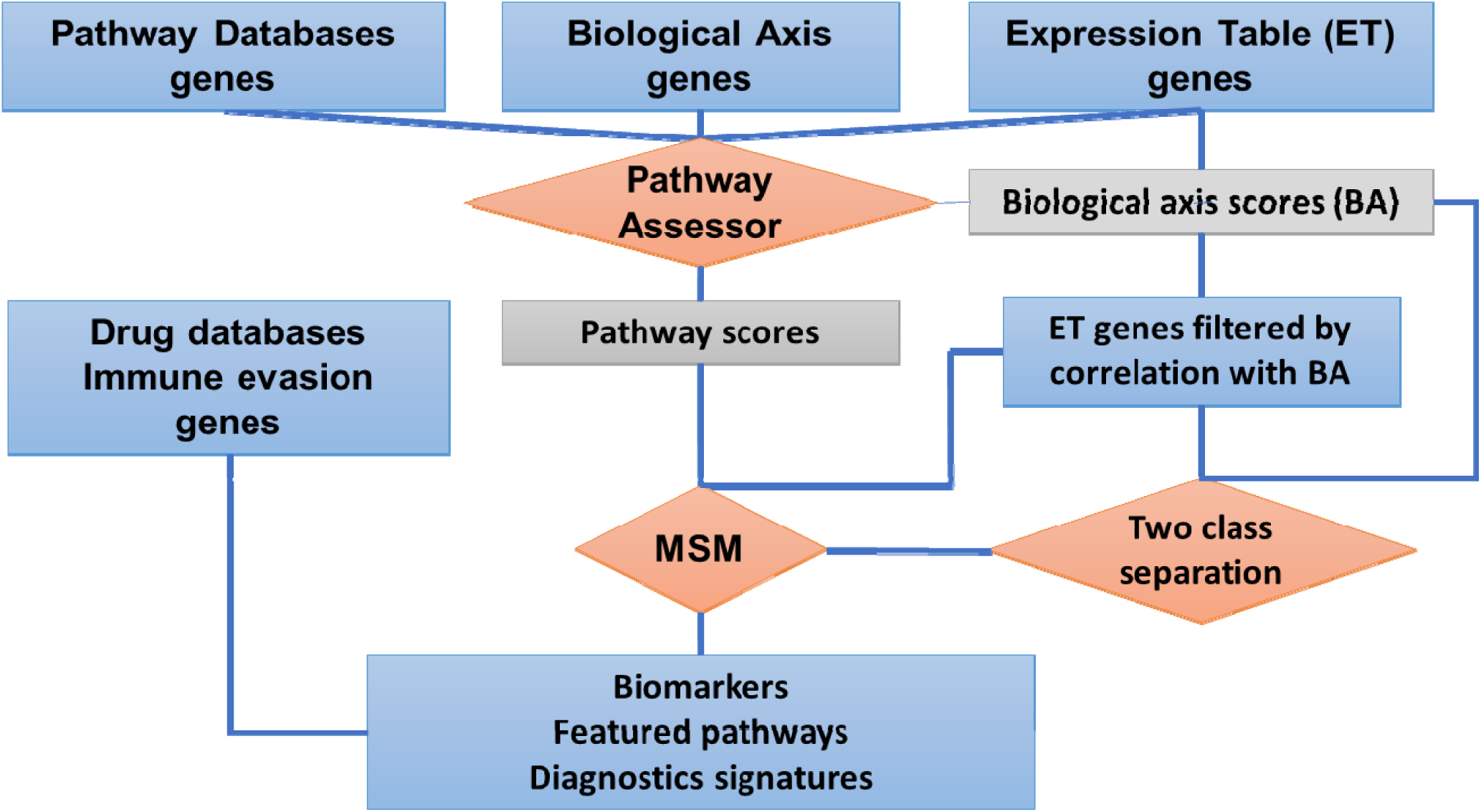
Computational protocol for classification of tumors into clinically distinct subtypes along a given biological axis. The protocol combines three algorithms (red diamonds): “Two class separation” (Supplement 1); the pathway scoring method “Pathway Assessor” (ssGSEA ^64^ and IPAS, Supplement 2); and the classification algorithm (Molecular Signature Method, MSM ^63^). Given as input gene sets (blue rectangles) from “Pathway Databases”, “Biological Axis”, “Expression Table” (ET), the “Pathway Assessor” computes scores (grey rectangles) “Pathway scores” and “Biological Axis scores” used for ranking tumors along this axis. “Biological Axis scores” can be input also as independent variable. The ET genes are been filtered by a given correlation (both positive and negative) with the BA scores, and, after this, given filtered the ET and the BA scores used by the “Two class separation” algorithm that finds the most distinct classes. Finally, the MSM ^63^ takes as input either the ET genes or Pathway scores and determines characteristic genes and pathways and their classification signatures. The obtained characteristic genes and pathways are been matched with genes available from genomic analysis of clinical data. In application to immune therapy, we use known genes and pathways involved in immune evasion.

In this approach, we assume that a “true” biological axis has to have the highest number of genes correlated with this axis; on the contrary, a biologically irrelevant axis correlates with any of the genes only by random. Therefore, for each of axes, we computed distributions of Pearson correlations between the axis and gene expression levels. In computational analysis we used gene expression levels of primary tumors of TCGA ^2^ and metastatic tumors of MSKCC ^13^. We tested three genes sets as clinically relevant axes and considered three types of pathway scoring. For nomination prostate tumors for PD-L1 therapy, we took the expression level of CD274 gene and inferred activities of the “IFN-γ/PD-L1 axis” pathway (Fig.1). This pathway is comprised of 29 genes, which can be classified as “positive” and “negative” regulators of CD274 expression. Assessing activity of the pathway, we used separately a subset of 17 positive regulators and combination of both positive and negative regulator genes (Fig.1).

The simplest way of scoring pathway activities is to compute a normalized sum of expression levels of all pathway genes; in more sophisticated approach – single sample gene set enrichment analysis (ssGSEA ^64^), one computes an enrichment score by assessing a shift of the actual distribution of pathway genes from the uniform distribution towards top (or the bottom) in the ordered gene list levels ^64^. In both of the approaches it is assumed that gene’s role in a pathway is entirely depends on its expression level; no specific context of expression levels of other genes or activities of molecular pathways is taken into account. One of the problems in such assessments is a potential presence of two brunches of genes in a pathway – positive and negative regulators. In our current tests of the pathway scores, we simply took the difference between scores of two brunches. To infer pathway activation more accurately, we introduced an entropy based assessment of deviation from uniform distribution our new entropy based score – IPAS (Inferred Pathway Activity score, Supplement 1). The main idea of the approach is to assess deviation from uniform distribution by computing two probabilities: one probability assesses enrichment of pathway genes at the top of ordered list of gene expression levels, the other one – from the bottom. Both probabilities can be significant, in case that a pathway has both significantly upregulated and significantly downregulated genes. The other essential difference is that a pathway can be assessed with a high scoring value even when only a few of pathway genes are significantly above the median of gene distribution, while the majority of pathways genes can be below the median. Thus, our scoring method is genuinely biased towards those pathways which have truly top (or bottom) regulated genes. In this study, we tested all above pathway scoring methods to choose the most accurate method for ranking prostate cancer tumors by inferred sensitivity to PD-L1 therapy.

The important part of the computation protocol is selection of characteristic genes and pathways and derivation of classification signatures. To this end, we used our previously developed Molecular Signature Method ^63^ and software.

## 4. Discussion and Conclusion

In this work, we hypothesized that response to PD-L1 checkpoint therapy depends primarily on activation of IFN-γ/PD-L1 signaling pathway (IFNG axis). This hypothesis means that non-responder tumors are either tumors with downregulated IFNG axis genes, which became invisible to immune cells, or tumors with upregulated IFNG axis genes and specifically activated mechanisms of immune evasion that prevent cancer cell death.

To test this hypothesis, we ranked tumors by computationally assessed activity of IFNG axis and assessed the biological relevance of the axis computing correlations between gene expression levels and the IFNG axis score. The more genes correlate with the axis, the deeper biological separation between tumors aligned along the axis. We also used the IFNG axis scores to nominate potential disease subtypes by determining a point of the maximal difference between two tumor sets with lower and higher levels of the IFNG axis score. Thus, we classified primary and metastatic prostate tumors into “IFN-γ-depleted” and “IFN-γ-enriched” subtypes along inferred activity of IFN-γ/PD-L1 signaling axis and determined genes and pathways differentially expressed and activated between “IFN-γ depleted” and “IFN-γ enriched” tumors. We examined four scoring of IFNG axis, including axis of PD-L1 (CD274) expression level alone and a new scoring function, IPAS, introduced for assessment of activation of molecular pathways. Based on correlation assessments, we concluded that taken alone an expression level of CD274 is not likely be a correct marker for immune therapy, and this is especially true for metastatic tumors. We settled on combination of IPAS scores computed for positive and negative regulators of IFN-γ signaling as the scoring axis, which produced the deepest separation of tumors into biologically different disease subtypes.

We obtained a number of positive controls in proposing IFNG signaling axis as clinically relevant. In particular, we found significant correlation between IFNG axis and immune system related pathways of all major pathway databases used ^65-69^, we also found significant correlation between the axis score and well known immune evasion genes (IDO1, ENTPD1). The genomic signatures showed significant prevalence of immunosuppressive cells over cytotoxic lymphocytes in “IFN-γ-enriched” tumors.

The consistency of results obtained for primary set and independently for metastatic set of prostate tumors suggest that overall, the revealed characteristic genes and pathways represent true biological differences between “IFN-γ-depleted” and “IFN-γ-enriched” subtypes of prostate cancer tumors.

The further analysis suggests optimization of weights of positive and negative regulators in IFNG axis scoring function, adding mutation burden as a co-factor in ranking tumors, identifying potentially missed regulator genes to the genes of IFNG axis – all these are new questions set by this study.

The obtained results set a problem of practical classification of prostate cancer tumors into “IFN-γ-enriched” and immune “IFN-γ-depleted” subtypes. The fact that big cohorts of prostate cancer tumors can be separated into different biological subtypes does not mean that such separation can be easily made for any new tumor expression profile. We proposed a solution to this problem, a biomarker panel composed of two gene sets, which are inversely regulated in IFN-γ-enriched and in IFN-γ-depleted subtypes. The IPAS scoring function can naturally combined two scores and produce an accurate ranking of tumors.

Thus, this study provides a basis for a rational design of novel quantitative platform for classification of immune phenotypes of prostate cancer tumors and development of combination therapy strategies for patients. It worth noting also that proposed approach can be applied not only to classification of immune phenotypes, but also to classification of tumors with known or hypothetic molecular mechanism of drug action, e.g. classification metabolic phenotypes along inferred activity of a drug specific metabolic pathway and also along experimentally derived drug sensitivity axis.

## Author contributions

BR, ES and AT designed the project; BR and ES developed the IPAS algorithm; BR and AC developed the software and performed data analysis; TO, BR, SN and AT performed biological and clinical interpretations of results; all wrote the paper.

## Data and materials availability

Supplementary:

Supplement 1. IPAS - a new method for comparison of tumors by Inferred Pathway Activation and Suppression Score.

Supplement 2. Algorithm for separation of tumors into 2 distinct classes along given scoring axis.

Supplement 3. Classification of different subtypes of primary and metastatic prostate tumors made with MSM algorithm.

Supplement 4. Statistical characteristics of genes and pathways differentially expressed between IFN-γ enriched and IFN-γ depleted subtypes (excel table).

Computational research reported in this paper was supported by the Office of Research Infrastructure of the National Institutes of Health under award numbers S10OD018522 and S10OD026880. The content is solely the responsibility of the authors and does not necessarily represent the official views of the National Institutes of Health.

## Supplement 1. IPAS - a new method for comparison of tumors by Inferred Pathway Activation / Suppression Score

Understanding the activation status of molecular pathways in tumors can help nominate disease driver pathways, identify targets for therapeutic intervention, and determine clinically relevant subtypes and diagnostic biomarkers. We introduce a new method to infer pathway activation and suppression by examining under- and over-representation of pathway genes in tumor genes ranked by gene expression levels.

Like in related ssGSEA method [1], we assume that biological activity of a pathway is regulated by collective changes of expression levels of majority of genes involved in this pathway; then a difference in a pathway activity between tumors can be assessed by a difference in positioning of expression levels of genes involved in this pathway in ranked list of expression levels of all genes in each of tumors (Fig.1).

The novelty of our approach rests in defining the pathway scoring function as an entropy difference associated with deviation of the gene expression levels from equilibrium or random distribution because of either activation or suppression. Following this idea, the entropy difference or scoring function can be presented as a log of ratio of two probabilities: a probability to occupy by random the observed positions in a list of tumor genes ranked by expression levels from the top to the bottom and, similarly, a probability to occupy by random the observed positions in a list of expression levels ranked from the bottom to the top. Thus, for a pathway of a single gene, its relative activity across tumors is defined as a negative log of ratio of two numbers: a number of genes with expression level bigger than an expression level of given gene, and a number of genes with expression levels less than an expression level of given gene.

**Fig.1.**
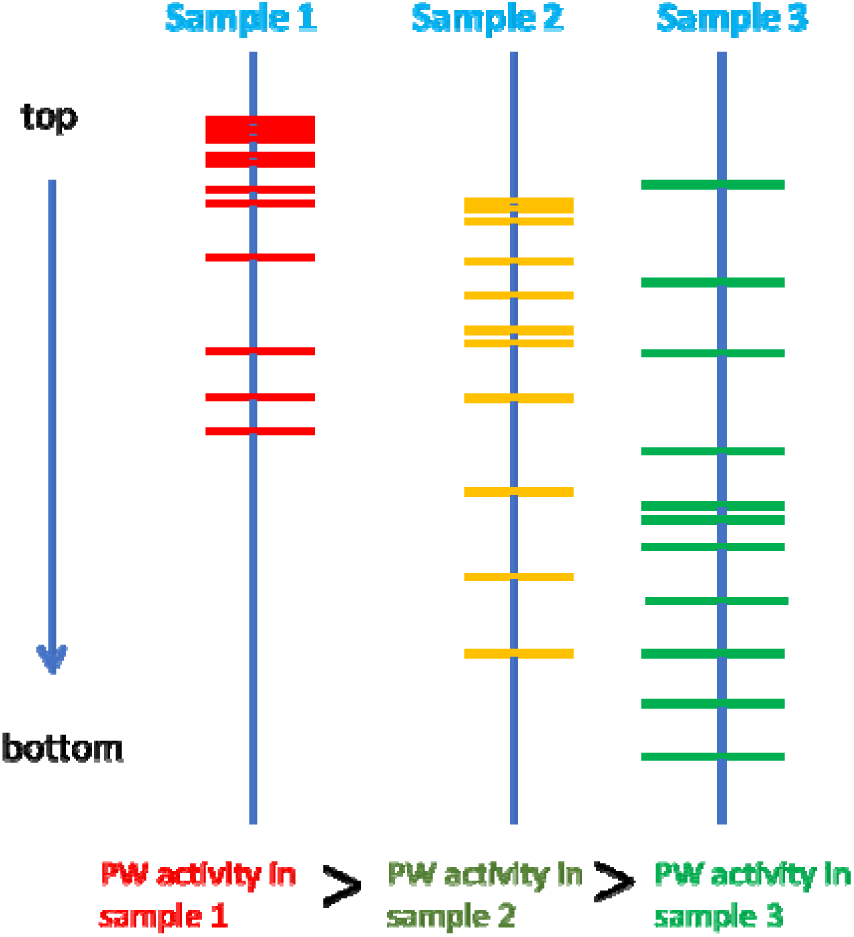
Tumors differ by distributions of gene expression levels. Expression levels of an illustrative pathway in different samples are shown by red, yellow and green bars. We hypothesize that activity of a given molecular PW in different tumors can be compared by levels of overrepresentation of pathways genes among top upregulated genes in a tumor. The roles and expression levels of individual genes are not taken into account in this approach.

For pathways of multiple genes, the “top” and “bottom” probabilities can be computed accurately by using statistical mechanics of one-dimension systems [2]. The complexity of this algorithms is assessed as ~M*N^2^, where M is a number of genes in a pathway and N is a number of genes in expression table. However, direct accurate computations of partition function become time consuming and impractical for M~200, N ~20K and ~5K pathways of major pathway databases.

Therefore, we assessed probabilities as geometrically averaged P values computed for each of genes using Fisher’s exact test [3], given gene’s ranks in a list of pathway genes and in a list of ranked genes of a tumor, a number of genes in a pathway, and the total number of genes with the assessed expression level in a given tumor (Fig.2).

**Fig.2.**
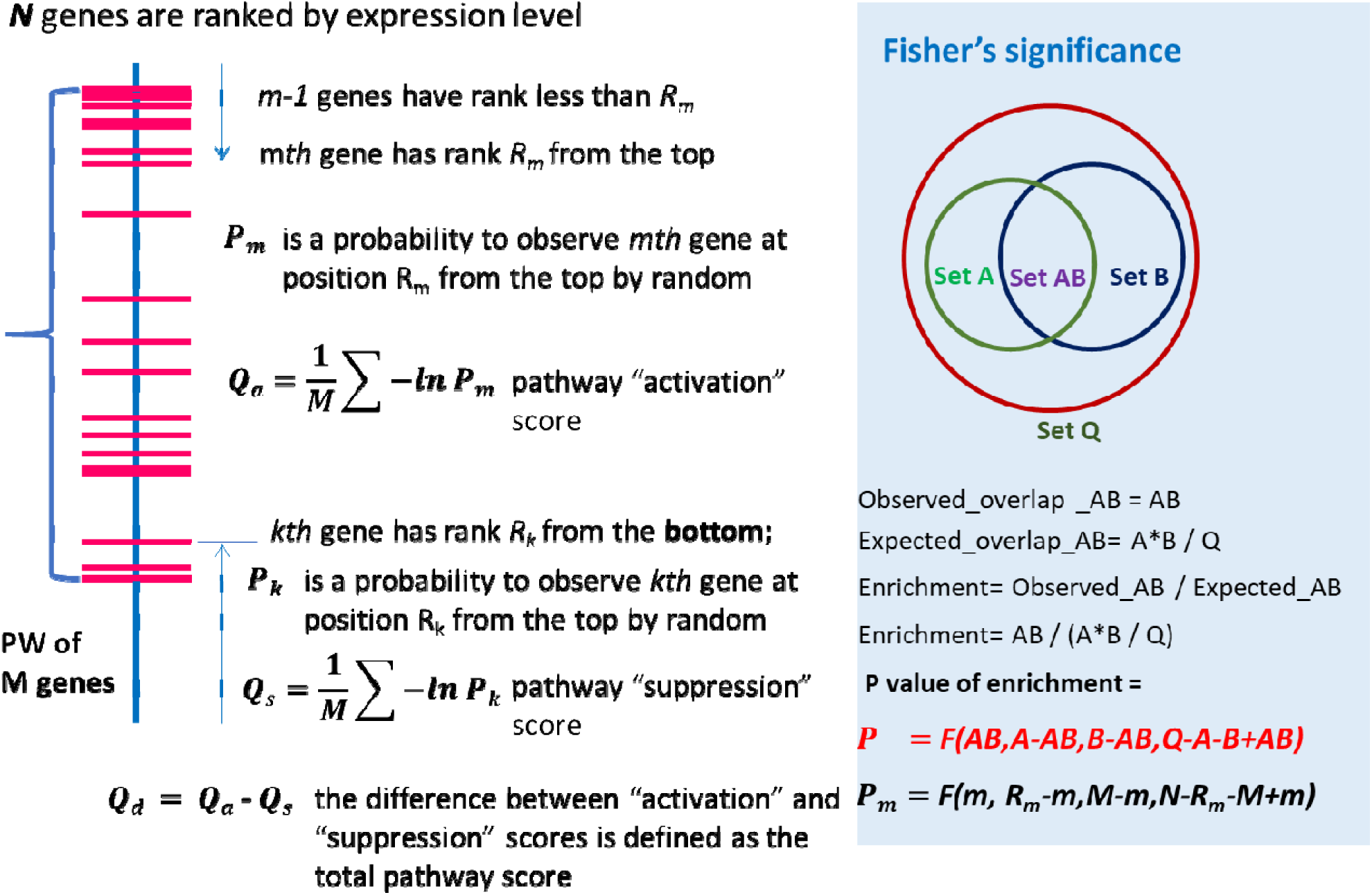
Algorithm for computing IPAS scoring function.

The scoring function *Q*_*d*_ is positive, when expression levels of pathway’s genes are overrepresented among top expressed genes of a tumor, and it is negative, when pathway’s genes are at the bottom of expressed genes of a tumor; the scoring function is close to zero, when expression levels are distributed by random or approximately equally split between genes at the top and genes at the bottom. Thus, considering independently activation, *Q*_*a*_, and suppression, *Q*_*s*_, scores, this approach makes possible to detect dysregulated pathways, which have gene expression levels overrepresented both at the top and at the bottom of a ranked gene list. Given a pathway with both positively and negatively regulated branches (e.g. cancer driver module which becomes active because both of upregulated expression of an oncogene and downregulation expression of tumor suppressor) one can assess its overall activity by combining inferred activities (or suppressions) of positive and negative pathway branches.

### Software and data access

To facilitate IPAS analysis, we have developed a web application accessible at: http://pathwayassessor.org. This tool allows researchers to perform IPAS analysis on a given expression table, without the need to install applications or write code. The user uploads an expression table, where row names are gene symbols and column names are sample identifiers. The user then selects several parameters, including: Pathway Database, Mode, and Direction. The final output is a table, or collection of tables, where the row names are pathway names, column names are sample identifiers, and each record is the final IPAS score.

The Pathway Database parameter determines the collection of pathways or gene sets for which to calculate IPAS scores. Pathway databases included are KEGG (Kyoto Encyclopedia of Genes and Genomes), Reactome, Hallmark gene sets from the Broad Institute, and HMDB/SMPDB (Human Metabolome Database and Small Molecule Pathway Database).

The Mode parameter determines the method of aggregating p-value scores for gene rankings to produce the final score. A user selects harmonic averaging, geometric averaging, or selection of the minimum p-value.

Direction options include “ascending”, “descending”, and “difference”. The options for “ascending” and “descending” indicate the sorting direction of a sample’s expression profile for determining gene rankings, for assessing the underactivity and overactivity respectively of genes in a pathway. The “difference” option produces a third table of scores with the difference between the overactivity and underactivity of genes in a given pathway.

When submitting the expression table and selected parameters, the user also enters their email address to receive analysis results. The analysis is then queued for execution on a remote server, which emails the results to the user as TSV format file attachments.

The source code for this web application is completely open-source and can be accessed on Github at: https://github.com/RevaLab/PathwayAssessor. For researchers who would like to include this analysis as part of a computational pipeline, the code is also available as a Python package, available through the Python Package Index (pypi) at: https://pypi.org/project/pathway-assessor/.

## Supplement 2. Algorithm for separation of tumors into two distinct classes along given scoring axis

Tumors ranked by a given scoring function. For a set of N tumors, there are N-1 possible separations into two classes (Fig.1). Which of those tumor classes are the most distinct?

Let *M* be a number of genes in a dataset; for a separation k, a difference between distribution of gene expression values in classes 0 and 1 can be assessed by a P value, P_km_. Then the statistical difference between gene expression distributions in two given classes of k and N-k tumors is assessed by a harmonically averaged P-value of individual genes:

**Fig.1.**
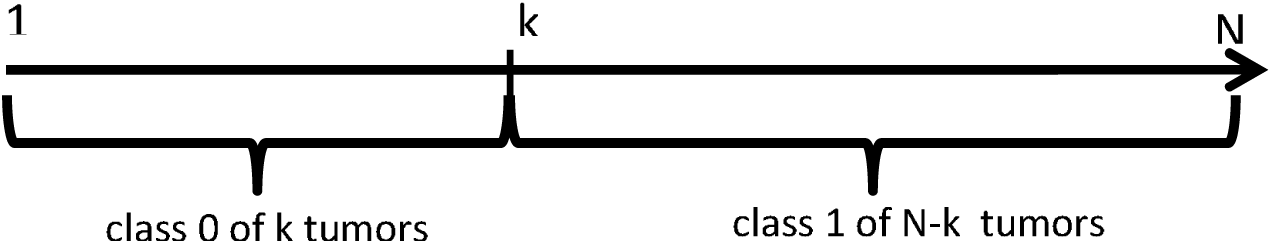
Separation of tumors into 2 classes along a given scoring function; there are N-1 separations into 2 classes.

It is assumed that the minimal value of determines the most distinct separation of tumors along a given axis. For control, we also performed random selection of tumors in classes 0 and 1, computed, {*P*_*mrk*_} and assess a random separation of tumors by harmonically averaged P-values {*P*_*rk*_*}*, where r =1, 2,… 10^4^.

For computing *P*_*mrk*_, we implemented and tested several methods. The most robust method, which never gives P=0 for practical gene expression distribution is describe below in Fig.2.

**Fig.2.**
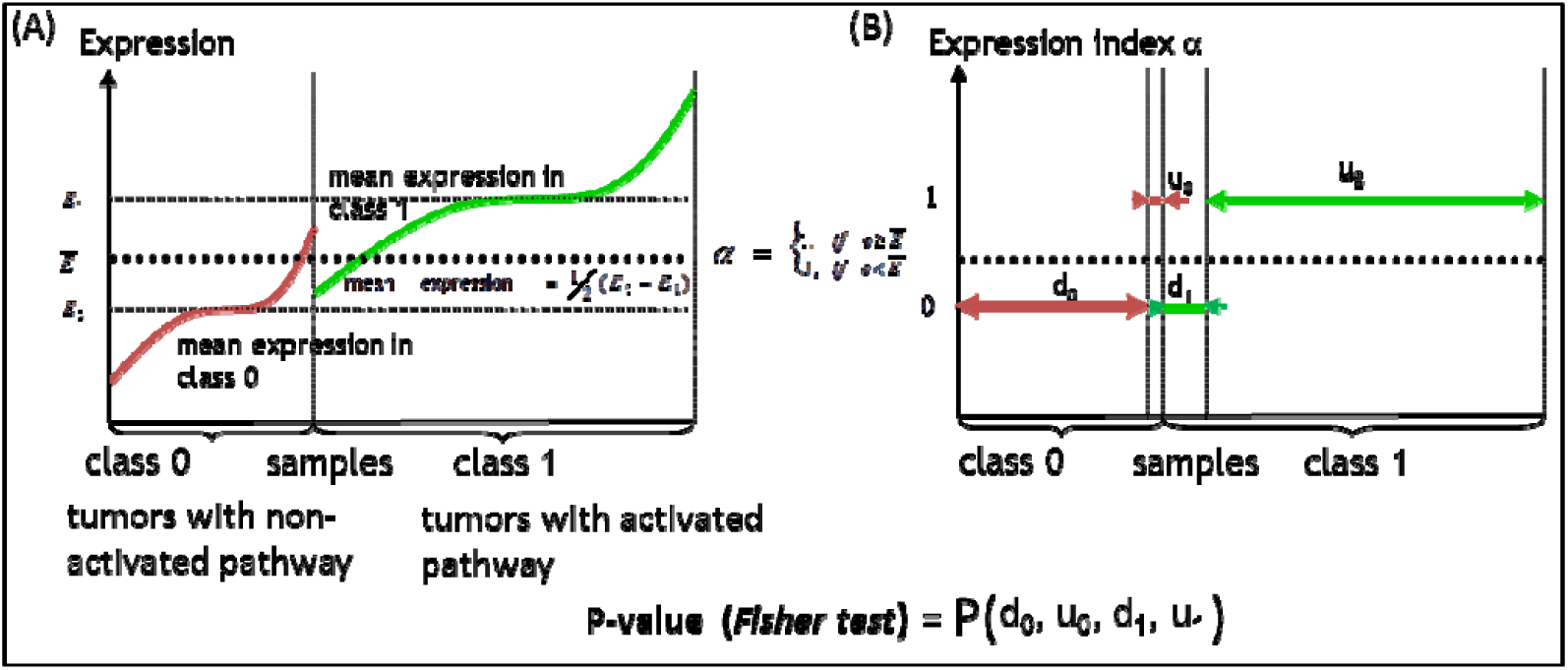
Computing a P-value between two distributions of gene expressions using the discrete approximation. (A) For each gene, the mean expression values, *E*_0_ and *E*_1_, are computed for sample classes 0 and 1, respectively. The discrete approximation threshold, *Ē*, is defined as the average value of *E*_0_ and *E*_1_. (B) A discrete index “u” or “1” is assigned to an expression value *e*, if *e* ≥ *Ē*; an index “d” or “0” is assigned to *e*, if *e* < *Ē*. Thus, depending on a gene’s expression values, samples of class 0 are separated into two groups, d_0_ and u_0_, and samples of class 1 are separated into groups d_1_ and u_1_, respectively. For clarity, expression values are sorted within each of the sample classes. The statistical significance of association between expression values and tumor classes is assessed by Fisher test [1].

## Supplement 3. Classification of different subtypes of primary and metastatic prostate tumors made with MSM algorithm

Given a table with gene expression levels and two tumor classes, the algorithm ^1^ uses halve of tumor samples as a training set to determine (i) biomarker genes, (ii) a weighted combination of biomarker expression levels (classification score) and (iii) optimal threshold for separation tumors of training set into given classes. Then, the determined biomarker genes, their weights and threshold were used in classification of remaining halve of tumor samples. After 1024 rounds of training/test classifications, for each of samples its probability of been correctly classified in training and test set was determined as well as averaged value of classification function. The classification accuracies were assessed by area under receiver-operating curve (AUC). Two AUC values reported in figures for both training and test classifications ^1^ were derived from, respectively, probabilities of each of samples to be classified in correct class; and from the average classification scores.

The figures below show classification accuracies of different disease subtypes computed for gene expression values of individual genes and for pathways scores; probabilities and averaged values of scoring functions are shown only for classification of test sets; red and blue circles represent tumors of different classes.

The presented classification plots show that revealed immune subtypes (Fig.S3.1, FigS3.3) are distinct as much as genomic subtypes in primary tumors (Fig.S3.4) or clinical subtypes in metastatic tumors (Figs.S3.5-8). They also show that the immune subtypes (Fig.S3.1, FigS3.3) can be classified with using relative expression values of a small number of biomarker genes (Fig.S3.2 and Fig.S3.4) practically as accurate as with using gene expression values or pathways scores assessed from ~20K expression values for each of the samples. Finally, we demonstrated the competiveness of classification based on pathway scoring as compared to individual gene expression levels and competiveness of a new IPAS scoring method as compared to ssGSEA. The pathway scoring is based on relative position of genes expression levels and therefore can be potentially platform independent.

---

**Figure S3.1.**
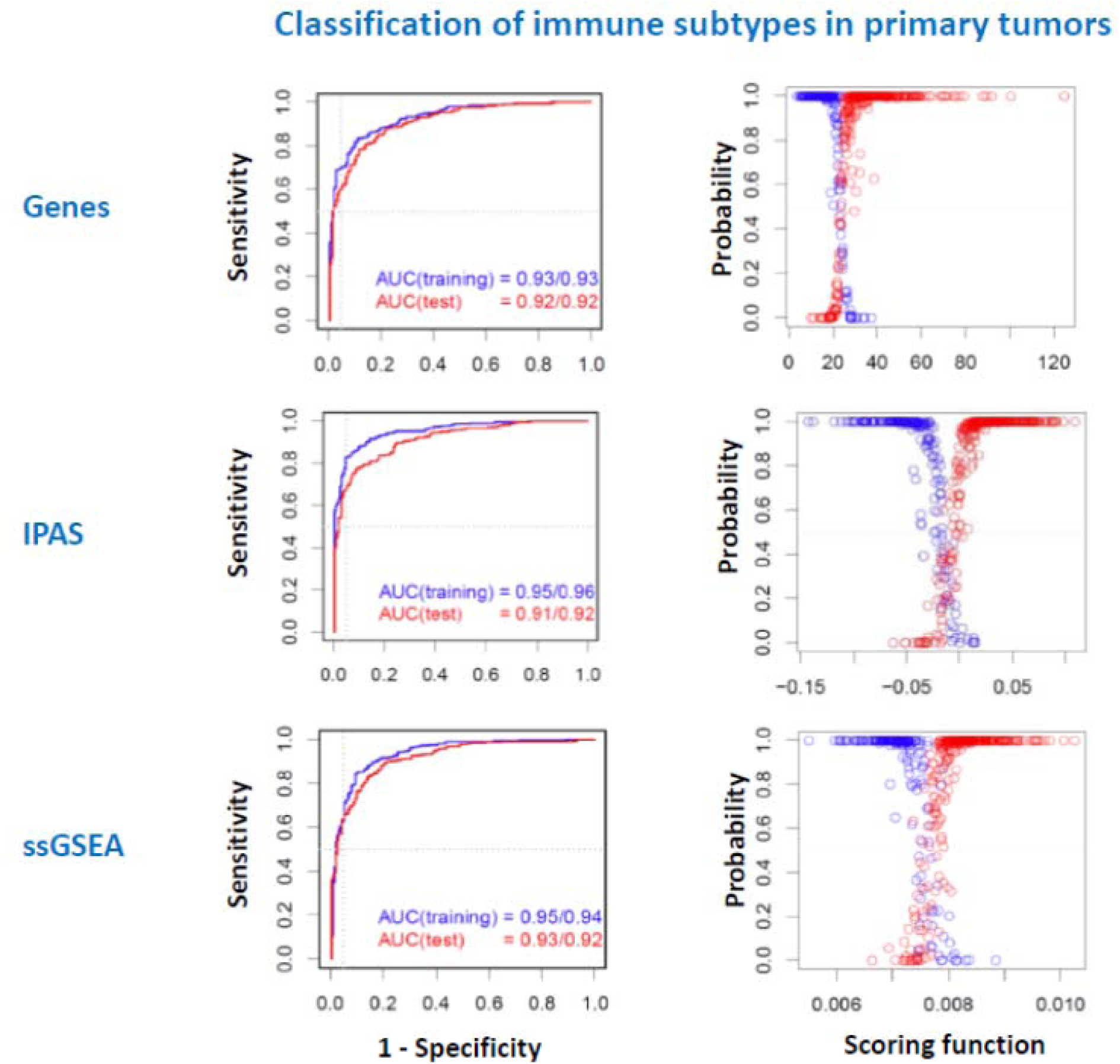

**Fig.S3.2.**
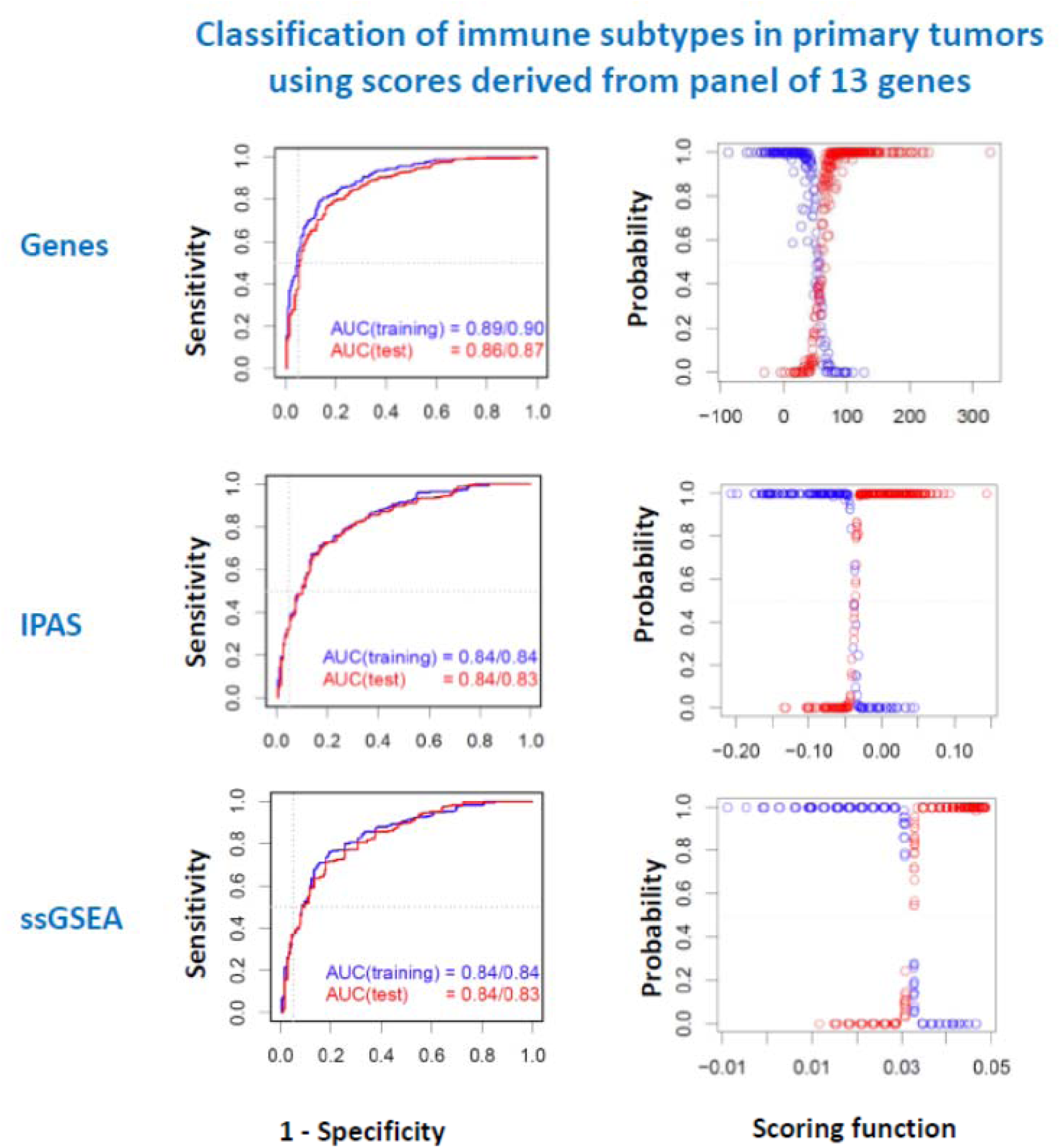

**Figure S3.3.**
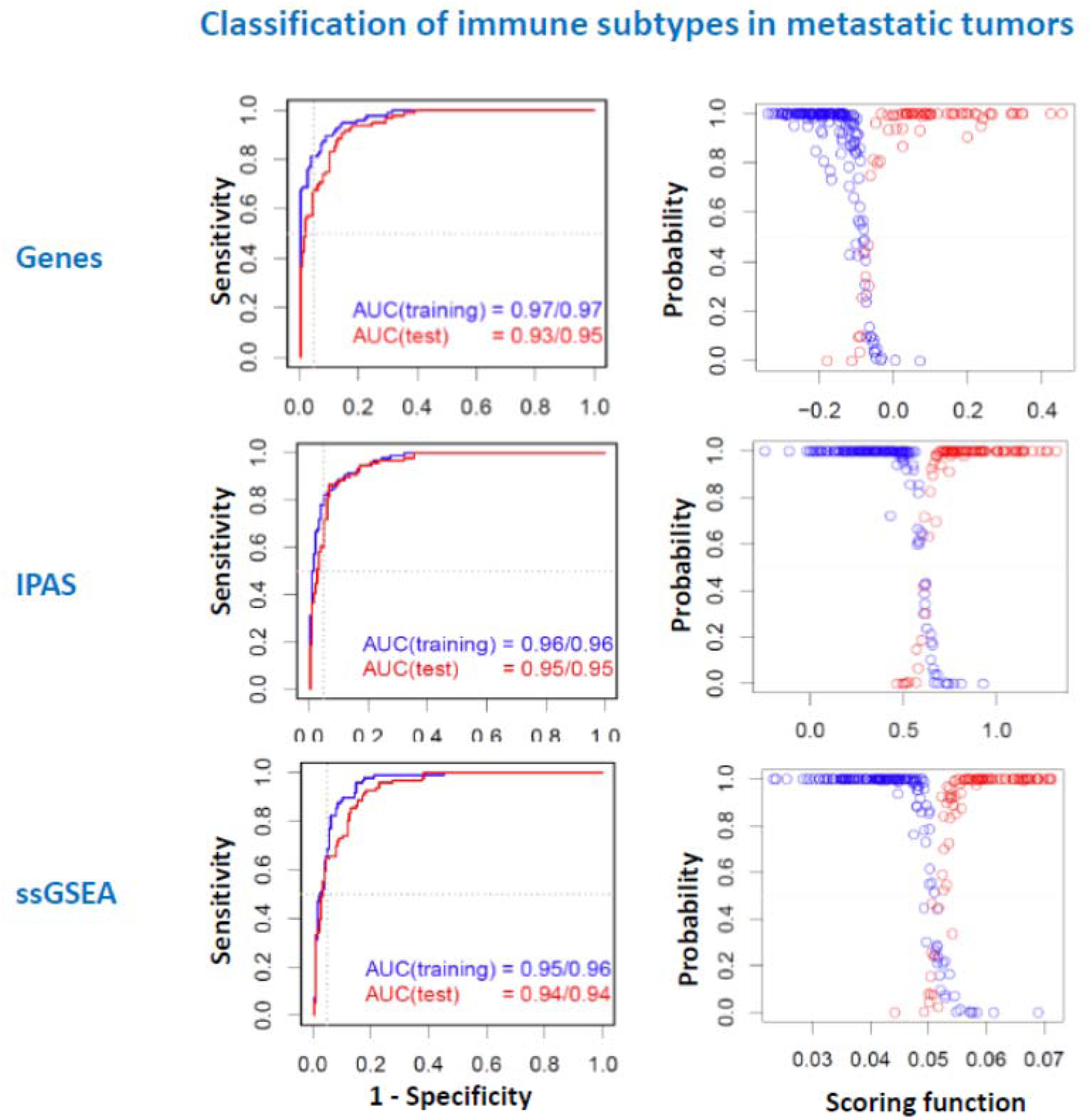

**Fig.S3.4.**
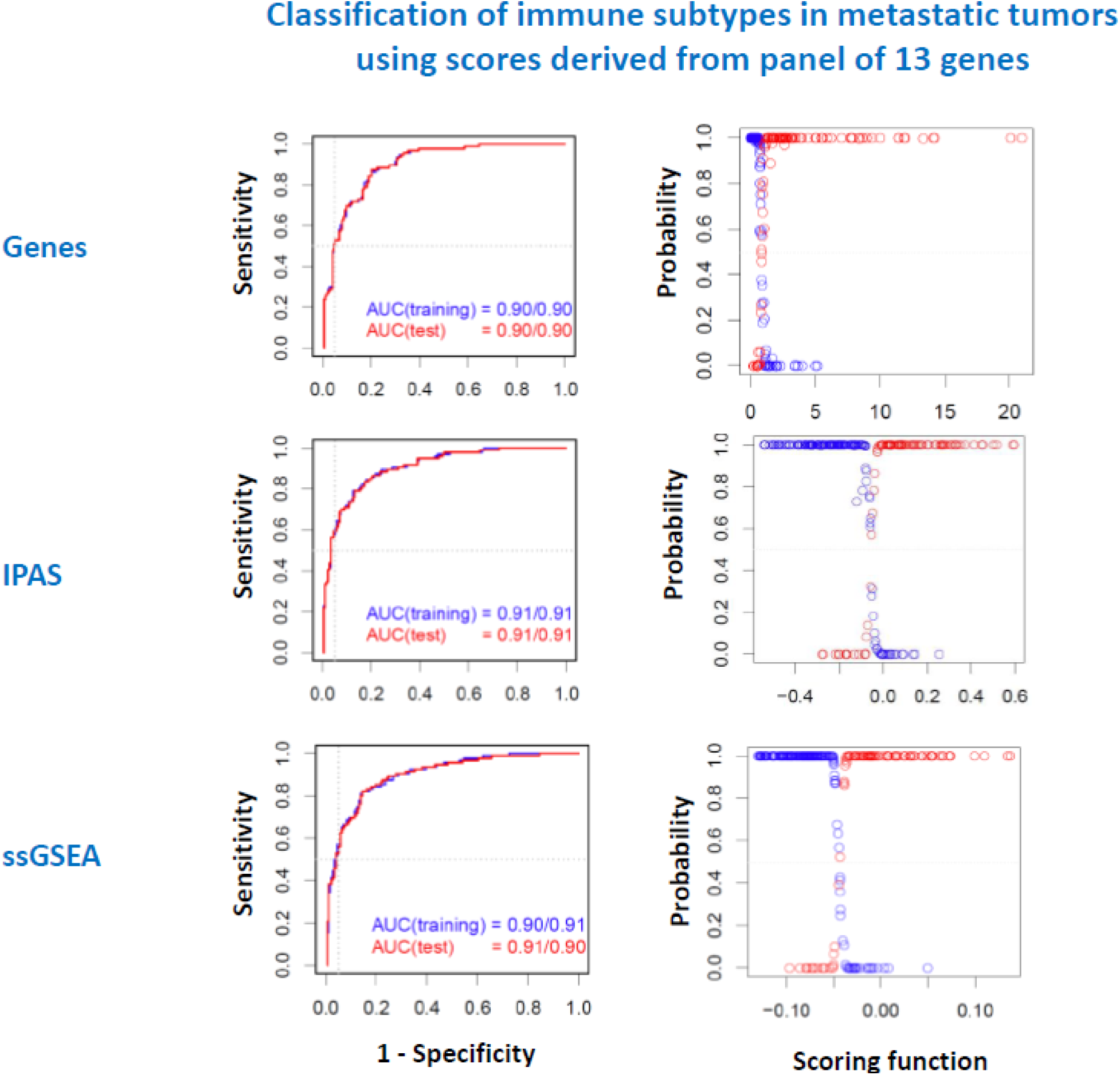

**Fig.S3.5.**
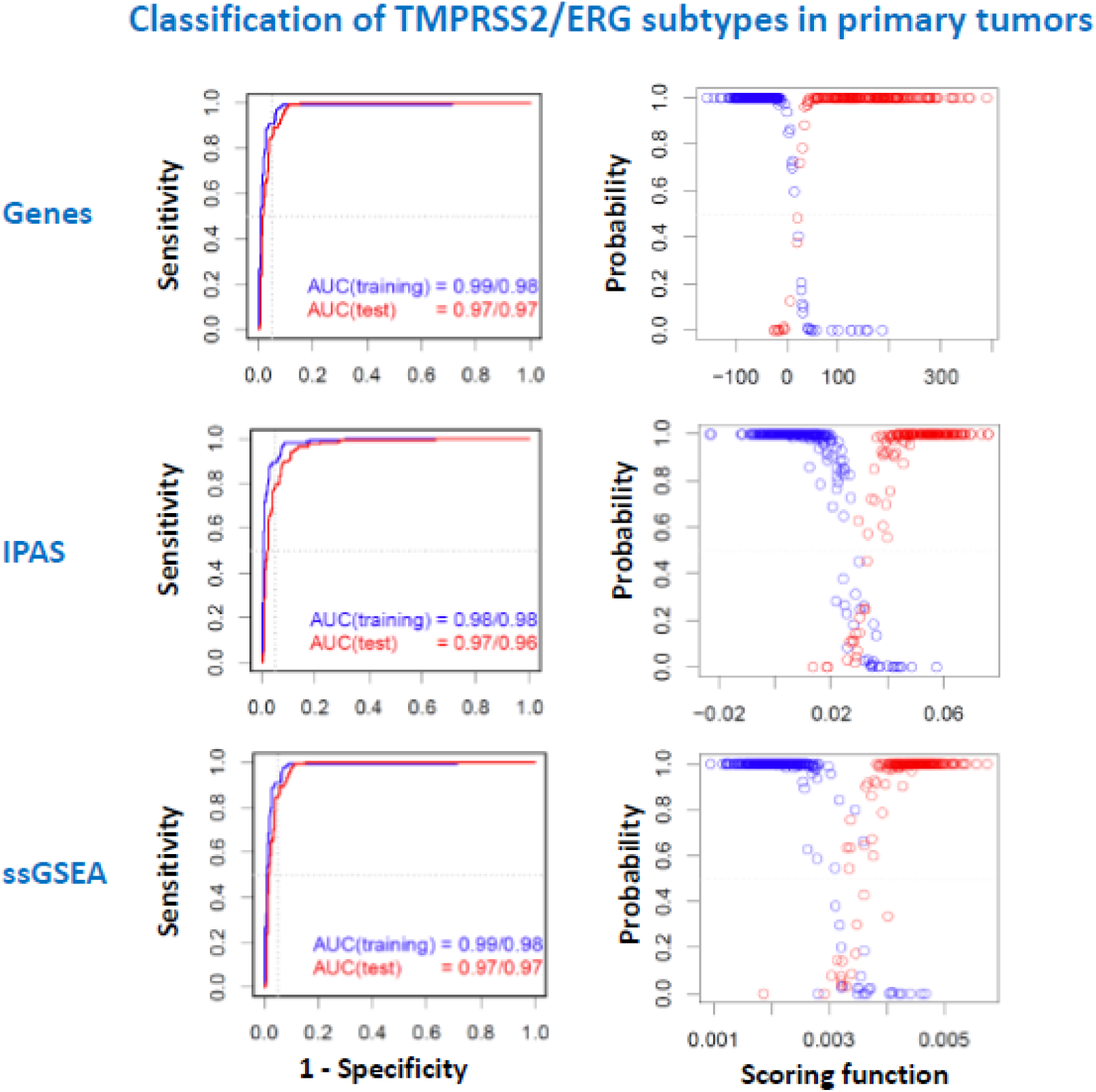

**Fig.S3.6.**
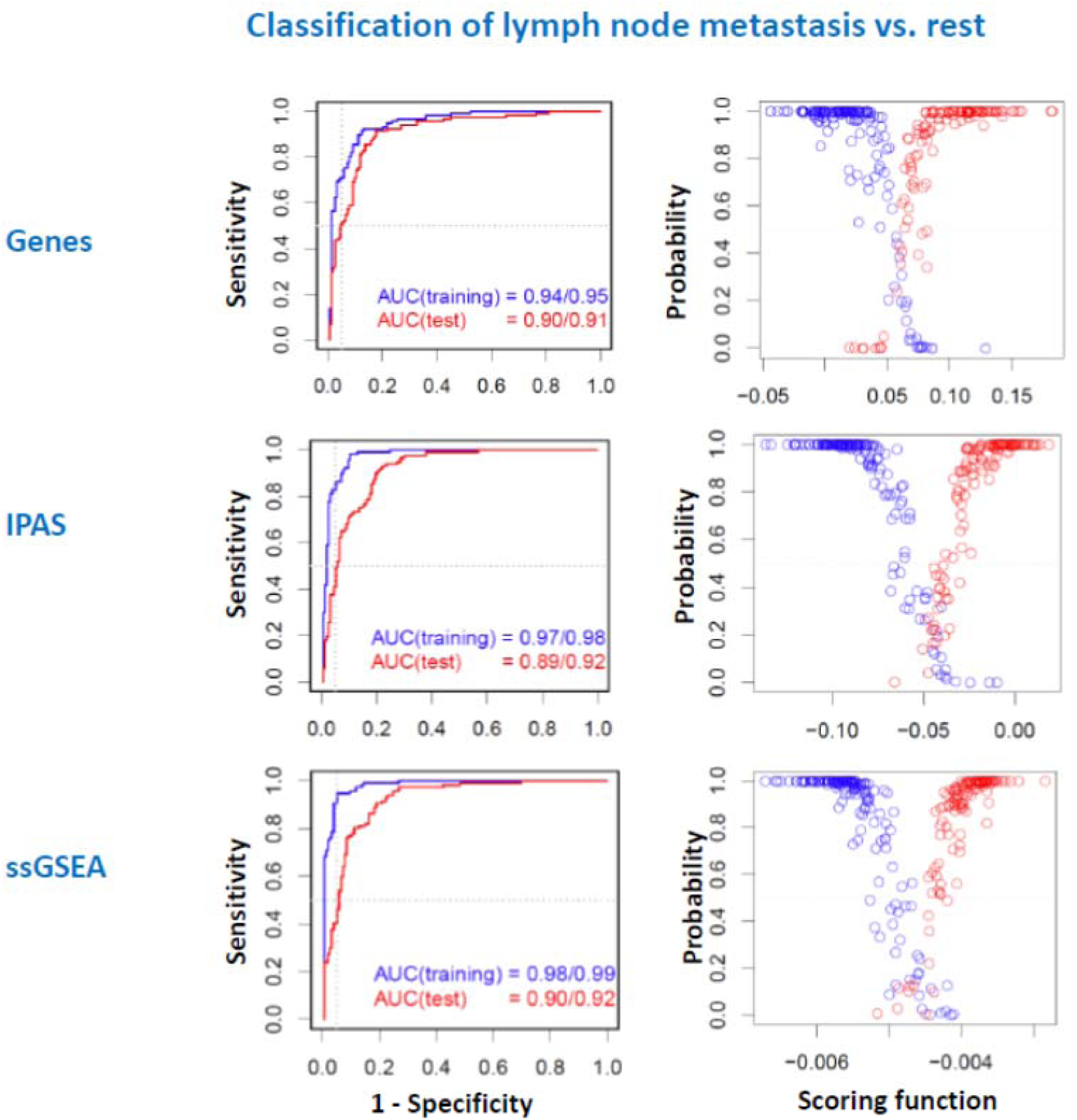

**Fig.S3.7.**
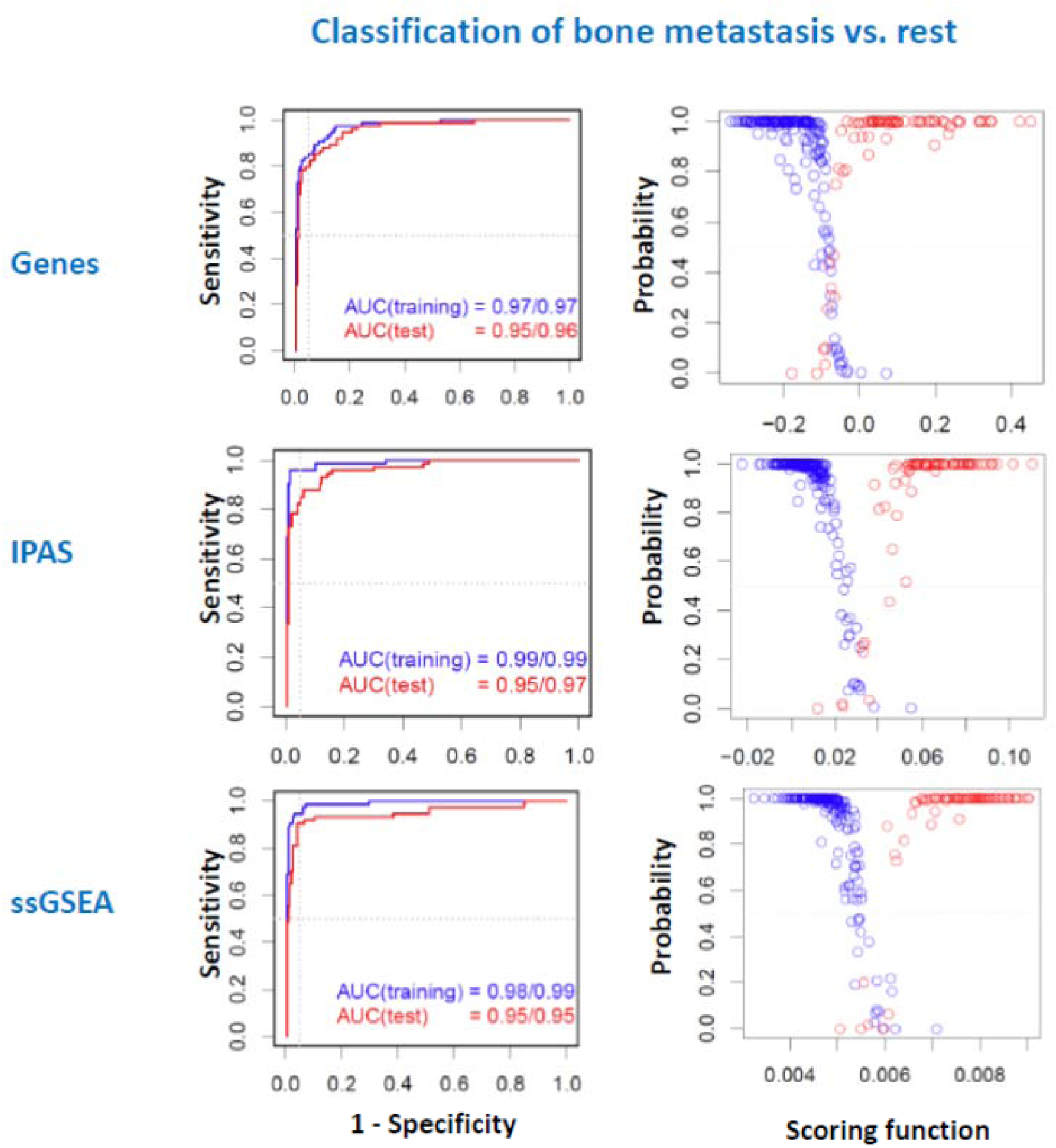

**Fig.S3.8.**
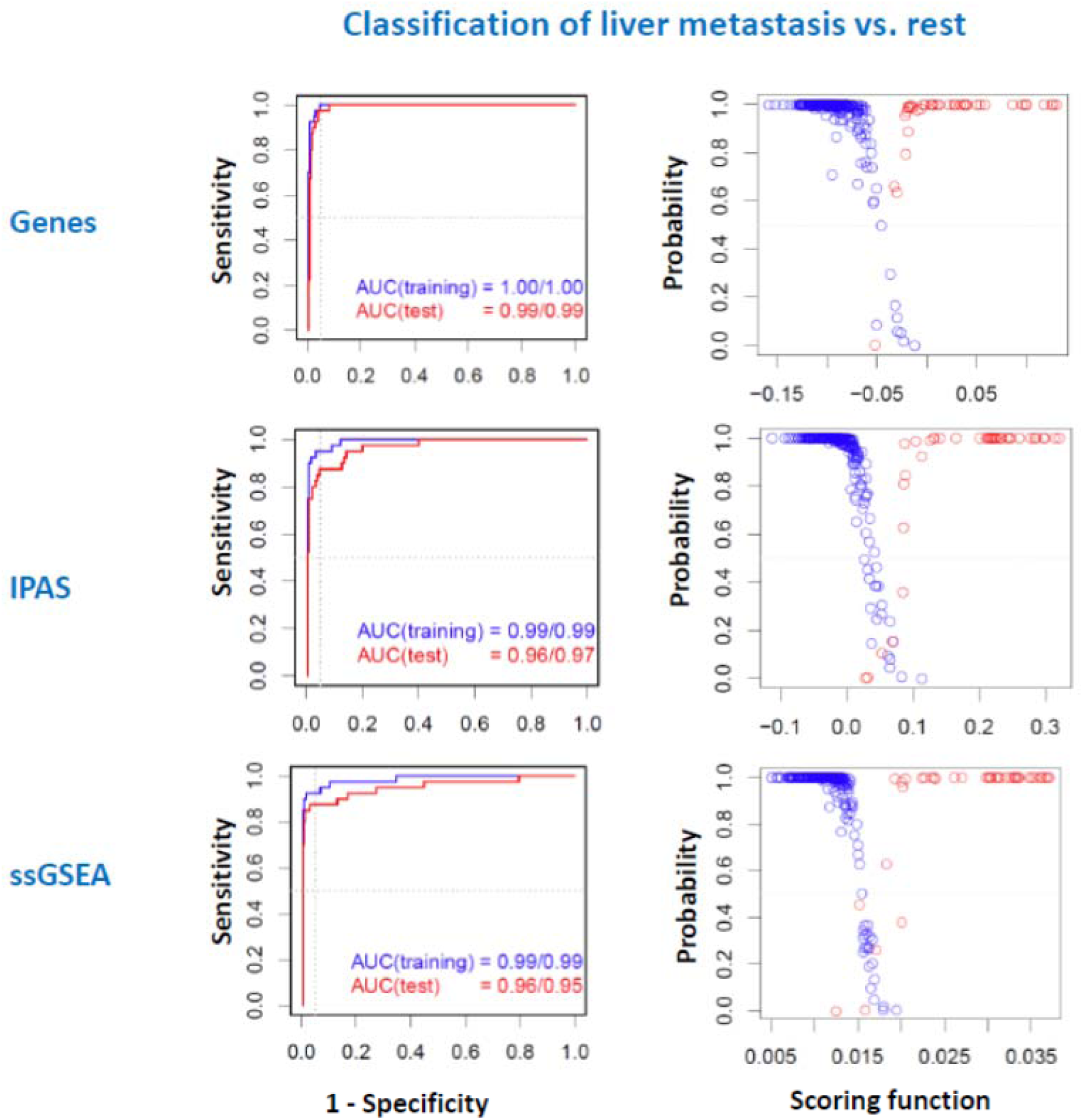

